# Wild genes to the rescue: High-throughput genomics reveals the wild source of broomrape resistance in sunflower

**DOI:** 10.1101/2025.09.14.676089

**Authors:** Dana Sisou, Hammam Ziadna, Mika Eizenberg-Weiss, Hanan Eizenberg, Sariel Hübner

## Abstract

- The ongoing evolutionary arms race between crop plants and their parasites necessitates a constant exploration of new genetic resistance. Broomrape (*Orobanche cumana*), a devastating parasitic plant, presents a formidable challenge to sunflower production, yet the genetic mechanisms underlying host resistance are still largely unknown.
- To address this gap, we developed a high-throughput phenotyping platform to quantify root infestation in a highly diverse sunflower association mapping (SAM) population. Using a dual GWAS approach with both SNPs and k-mers, we were able to pinpoint the genetic basis of resistance.
- Our findings validate previously identified QTLs with greater resolution and reveal several novel candidate genes conferring resistance, including putative leucine-rich repeat receptor kinases. Critically, the k-mer mapping approach circumvented reference genome bias, highlighting key introgressions from wild *Helianthus* species that have contributed to broomrape resistance.
- This research provides a powerful methodology for gene discovery and demonstrates that wild relatives remain a vital source of genetic material, offering breeders a significant advantage in the ongoing battle against rapidly evolving parasites.

## Introduction

Parasitic weeds represent a significant global threat to crop production (Fernández-Martínez et al., 2015). Broomrapes (*Orobanche* and *Phelipanche* spp.) are holoparasites that lack the ability to perform photosynthesis, instead drawing all necessary nutrients directly from their host plants (Molinero-Ruiz et al., 2015). This parasitic relationship results in substantial reductions in crop yield (Eizenberg et al., 2004; Molinero-Ruiz et al., 2015; Parker, 2009). In particular, the sunflower broomrape (*Orobanche cumana Wallr*) is a major constraint to sunflower production across regions including the Middle East, Central and Southeast Europe, and parts of Asia and Africa (Duca et al., 2022). Although some chemical control treatments exist, the most effective, sustainable, and environmentally friendly strategy for reducing broomrape infection is through the development of resistant cultivars (Fernández-Martínez et al., 2015; Pérez-Vich et al., 2004).

The defensive mechanisms which provide resistance against broomrapes can operate at different stages of the parasitism in accordance with the parasitic life cycle: (I) pre-attachment mechanisms, occurs during the broomrape seed germination and attachment of the seedlings to the host root (Yoder & Scholes, 2010; Yoshida & Shirasu, 2012), (II) pre-haustorial mechanisms, in which the penetration and establishment within the host roots is halted by physical barriers such as protein cross-linking, callose depositions and lignification (Echevarría-Zomeño et al., 2006; Letousey et al., 2007; Pérez-de-Luque et al., 2006; Pérez-De-Luque et al., 2007; Sisou et al., 2021), and (III) post-haustorial mechanisms, which takes place after the establishment of vascular attachments in the roots and consist with sealing of xylem vessels and blocking the flux of water and nutrients from the host to the parasite, leading to necrosis and subsequent death of the tubercles (De Zélicourt et al., 2007; Letousey et al., 2007; Martín-Sanz et al., 2020). Genetic resistance to broomrape is primarily race-specific and vertical (Molinero-Ruiz et al., 2015; Vrânceanu A.V et al., 1980), namely a gene-for-gene mode of resistance with dominant genes denoted as *HaOr1-HaOr7* confer resistance to broomrape races A-G (Duriez et al., 2019; Imerovski et al., 2016; Lu et al., 1999, 2000; Tang et al., 2003). To date, only one major resistance gene to broomrape has been identified; the *HaOr7*, which has been mapped to chromosome 7 and encodes a leucine-rich repeat (LRR) receptor-like kinase (Duriez et al., 2019). Other QTLs linked to broomrape resistance were mapped over the years, revealing a more complex genetic control (Akhtouch et al., 2016; Calderón-González et al., 2023; Imerovski et al., 2019; Louarn et al., 2016; Pérez-Vich et al., 2004; Pubert et al., 2024). Despite this progress, resistance in sunflower was eventually broken by more virulent broomrape races, motivating scientists and breeders to search for new sources of genetic resistance. Traditional landraces and mainly wild relatives to sunflower have been identified as beneficial source of diversity, providing plant breeders a rich resource of genetic variation for enhancing biotic and abiotic stress resistance (Seiler & Jan, 2014). Indeed, several resistance genes to broomrape were introgressed into cultivated lines from wild relative to sunflower including *H. anomalus* and *H. debilis* (Calderón-González et al., 2024; Cvejić et al., 2020; Fernández-Aparicio et al., 2022; Seiler & Jan, 2014). For example, QTLs such as *Or*_*Deb2*_ and *Or*_*SII*_ that confer resistance to broomrape were mapped within the same region on chromosome 4, where LRR-receptor like proteins (RLPs) and receptor-like kinases (RLKs) are located (Fernández-Aparicio et al., 2022; Martín-Sanz et al., 2020). A recent study has successfully mapped the *Or*_*Anom1*_ resistance gene which was introgressed from *H. anomalus* also to chromosome 4 at a genomic region enriched with receptor-like kinase genes (Fernández-Melero et al., 2024).

Genome wide association studies have been broadly used for identifying disease resistance genes in different plant species (Demirjian et al., 2023). In sunflower, a diversity panel named the sunflower association mapping (SAM) population was developed to capture much of the diversity in cultivated sunflower (Mandel et al., 2011). This population was thoroughly genotyped using whole genome sequence data and has been studied extensively to identify genes of interest (Gao et al. 2019; Hübner et al., 2019; Mandel et al., 2013). Nevertheless, it is now clear that a genotyping procedure that relies on a single reference genome does not capture the full diversity in a gene pool, and tend to miss an important fraction of the dispensable genes of which many confer resistance to biotic stress and were introgressed from wild relatives (Huang et al., 2023; Hübner et al., 2019; Hübner 2022a). Recently, a pangenome GWAS approach has been incorporated yet its implementation in large and complex datasets like sunflower is still sparse. To overcome this, a *k*-mer (small subsequences of length *k*) approach was suggested to identify polymorphism that is not represented in a reference genome which can be used in a GWAS framework without exceeding a reasonable computing cost (Voichek & Weigel, 2020). Combining different GWAS approaches can potentially improve the detection power of new candidate genes that confer broomrape resistance.

In this study, we investigate the genetic basis of broomrape resistance in the SAM population using SNP and k-mer GWAS. We developed a high-throughput screening platform to phenotype the roots of 1,413 samples infested with two distinct broomrape races. Our results confirm the contribution of previously identified QTLs and highlight novel genes for broomrape resistance, many of which were introgressed into cultivated lines from wild sunflower relatives.

## Materials and Methods

### Plant material

The sunflower association mapping (SAM) population was used to study the susceptibility of cultivated sunflower to two broomrape races. The SAM population is comprised of 287 accessions including inbred lines, open-pollinated varieties and landraces, thus the collection captures most of the allelic diversity in cultivated sunflower (Mandel et al., 2011). For control, two Israeli varieties, DY3 and EMEK3 (kindly provided by Sha’ar Ha’amakim Seeds Ltd.) were included in the study as known susceptible and resistant lines, respectively.

Two populations of sunflower broomrape were used to inoculate the SAM population: ‘Yavor’ population, which was collected in an infected field during 2014 in Western Galilee; Israel (32°53’48.0”N 35°10’39.4”E), and the ‘Gadot’ population which was collected during 2019 in an infected field in Kibbutz Gadot, Upper Galilee (33°0’43.56”N, 35°36’30.2328”E). The two populations were reported to be distinct races, where the ‘Gadot’ race is more virulent to the local resistant variety than other broomrape populations in Israel (Eizenberg et al., 2004). Broomrape seeds were extracted from the capsules using a 300-mesh sieve and stored in the dark at 4°C. Seed germination was evaluated with 10^−5^M of the synthetic artificial germination stimulants GR24 and Dehydrocostus lactone (DCL) before use according to a standard protocol (Plakhine & Joel, 2010) for broomrape germination (Fig. S1).

### Evaluation of response to sunflower broomrape

One seed from each sunflower accession was germinated and grown until a root system had started to develop. Seven-day-old sunflower seedlings were germinated in sterilized sand to minimize pathogen contamination and transplanted onto glass-fiber filter paper (GF/A, Whatman International Ltd.) placed over 100 mL of vermiculite in 12 cm × 12 cm square petri dishes. The petri dishes were pre-drilled at the top and bottom to accommodate shoot growth and root water uptake (Fig. S2). Sunflower broomrape seeds were surface sterilized with 70% ethanol for 2.5 minutes, followed by 1% sodium hypochlorite for 10 minutes, and then rinsed five times with sterile distilled water. Approximately 30 mg of sterilized seeds were uniformly spread onto the GF/A paper containing the sunflower seedlings in each dish. The dishes were then placed in plastic containers within a controlled growth chamber (25°C, 18-hour light, 150-200 µE m^−2^ s^−1^), and half-strength Hoagland nutrient solution (Hoagland D.R & Arnon D.I, 1950) was provided as needed.

Thirty-five days post-infestation, the response of each sunflower accession was evaluated. Root systems in this rhizotron-plate system were scanned using a high-resolution scanner (EPSON 12000XL America Inc., USA), and the resulting images were analyzed with ImageJ (Schneider et al., 2012). For each plant, the number and area of broomrape tubercles, as well as the sunflower root area, were quantified. Tubercles were categorized as healthy or necrotic. The response was recorded as attachment density, defined as the number of attachments normalized to the root area (cm^2^), for healthy (HealAD), and necrotic (NecAD) attachments (Fig. S3). All plates were manually inspected to validate the image analysis scores and the number of broomrape attachments were corrected to the size of the root system. We used a generalized linear model (GLM) to compare phenotypic traits between the four main genetic clusters in the SAM population (HA-Oil, HA-NonOil, RHA-Oil, RHA-Non-Oil), followed by post-hoc analysis (multcomp v1.4.26,Hothorn et al., 2008) of the estimated marginal means (emmeans v1.10.5,Lenth, 2024).

### Field trials

Field trials were conducted during late spring-summer (May-August) at two locations in 2020 and 2021 respectively. The sunflower response to the ‘Yavor’ broomrape population was assessed during 2020 at “Gadash farm”, Upper Galilee, Israel (33°17’96.53”N, 35°58’33.75”E). The experiment followed a randomized block design with four replicates (blocks) for each treatment (infested/control), where each replicate unit was composed of a four-plants plot of the same accession. Plants were pre-germinated in trays and transplanted to a weed free field with no background of broomrape infestation, thus we artificially inoculated the ‘treatment’ blocks with a 50 ppm broomrape seed/soil admixture added to each planting pit. The second experiment was conducted in 2021 at a naturally infested field with ‘Gadot’ broomrape population near Kibbutz Gadot, Israel (33°01’16.8”N, 35°36’31.5”E). To reduce infestation in the control plots, half of the field was fumigated with 600 g/ha formulated metam-sodium (Edigan, 370 g a.i./L) 21 days before sunflower planting, following soil sterilization procedures (Goldwasser et al., 1995). To verify sterilization efficacy, the susceptible cultivar DY3 was sown along the perimeter of the control plot. The experiment was conducted in a randomized block design with four replicates for each treatment (infested/control), where a replicate unit was composed of 6 plants sown in a row with distance of 40cm between plants. In both field experiments the blocks were grown on 1.96 m wide beds, with each bed containing two rows and an in-row spacing of 40 cm between plants within a replicate unit, and 80 cm spacing between units (Fig. S4).

In the field experiments, the following phenotypic traits were recorded during the growing seasons: number of days to budding, days to anthesis, branching (number of flowers per plant), plant height (ground to the basis of the head), stem diameter, main flower diameter, total plant biomass, and the number of broomrape shoots that had emerged per sunflower plant. Phenotypic variances were estimated with a linear mixed model (LMM) having the accession, treatment (plots) and the interaction between them considered as fixed effects and block (replicate) as a random effect. LMM was followed by ANOVA using the lme4 (Bates et al., 2015) and car (Fox & Weisberg, 2019) R packages.

### Genotype data

Genotype data for the entire SAM population was obtained from the sunflower-genome repository (https://www.sunflowergenome.org/) based on an extensive variant calling procedure conducted for this collection using the HA412-HOv2 as reference genome (Todesco et al., 2020). The genotype dataset was filtered using a minimum minor allele frequency of 5%, and a maximum of 10% missing data before analyses wherever applicable.

In addition, we genotyped the entire SAM population for *k*-mer presence/absence variation using the kmerGWAS package (Voichek & Weigel, 2020). Briefly, a list of *k*-mers (*k*=31) was called directly from raw sequence data after filtering low quality reads using Trimmomatic v.036 (Simon Andrews, 2020). Lists were then processed to distinguish between canonized and non-canonized *k*-mers using KMC v3 (Kokot et al., 2017), and counts from each accession were combined to one conclusive table. The genotyped *k*-mers table was then filtered to include calls that appear in at least five accessions and have appeared in canonized/non-canonized form at least 20% of the accessions. Finally, a *k*-mer table containing the presence/absence variation (PAV) of each *k*-mer across the entire SAM population was obtained.

### Genome wide association analyses

All genome wide association analyses were performed with GEMMAv0.98 (Zhou & Stephens, 2012) using a linear mixed model (LMM) and correcting for population structure and relatedness among accessions. The analyses were conducted for phenotypic scores obtained from the rhizotron platform for both ‘Yavor’ and ‘Gadot’ races. To improve the detection power of QTLs, we adopted an extreme phenotype approach which was previously shown to be efficient for mapping complex diseases (Amanat et al., 2020; Li et al., 2019). Phenotypic scores were converted to a case/control setup using a stringent cutoff to define resistance (controls) across all replicates screened in the rhizotron, where the cutoff for susceptibility was determined for each trait based on its distribution across accessions (HealAD > 0.5, NecAD > 0.3; Fig. S3). Thus, accessions with complete resistance were scored as 0, susceptible accessions were scored as 1, and ambiguous accessions were excluded from the analysis per trait.

In each model, population structure was controlled using the first two PCs obtained for a principal component analysis (PCA) conducted in PLINKv1.90b6.4 (Chang et al., 2015). The SNP dataset was filtered at a minor allele frequency of 5% and maximum of 10% missing data before each analysis for consistency. Statistical inference was determined using the Wald test and p-values were corrected for multiple testing with the SimpleM algorithm (Gao, 2011) and Bonferroni correction at 5% threshold.

For the *k*-mer GWAS, we used GEMMA with default parameters after correcting for population structure and relatedness based on PCA and kinship calculated directly from the *k*-mer PAV matrix. A -log10 threshold for the 10% family-wise error rate was considered as significant *k*-mers associations with each trait. Significant *k*-mers from the GWAS analyses were mapped to the HA412-HOv2 reference genome, as well as to three additional sunflower cultivar genomes: the maintainer line XRQv2, the restorer line PCS8, and the introgression line LR1 (https://www.heliagene.org/; Huang et al., 2023). Mapping of *k*-mers to each reference genome was conducted using *bwa aln* tool in BWAv0.7.17 (H. Li & Durbin, 2009) and parameters were set to ensure zero gaps and missingness within *k*-mer (-o 0 -n 0). The alignment files were processed, indexed and sorted using samtools.v1.17 (Danecek et al., 2021), and only uniquely aligned reads (XT:A:U tag) with mapping quality > 20 were kept. Mapped *k*-mers were extracted and clustered by calculating pairwise LD between *k*-mers using the formula:

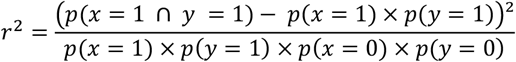

Where *p* denotes the probability (based on the observed frequencies) of two *k*-mers (*x* and *y*) to be present or absent (1 = presence, 0 = absence) separately or jointly (∩) across accessions. *k*-mers clusters were then ordered by the trait value (phenotypic response) of each accession and plotted using ggplot2 (Wilkinson, 2011).

### Identification of candidate genes, pathways and introgressions

To search for potential candidate genes around significant associations of SNPs and *k*-mers, we calculated the linkage disequilibrium (LD) decay in the SAM population with PopLDdecay (Zhang et al., 2019). The size of the regions to be explored was determined at ± 250Kbp based on the LD decay (r^2^ = 0.3; Fig S9) among SNPs. Lists of candidate genes were retrieved from the HA412-HO.v2 annotation file by determining the interval around significant SNPs and *k*-mers. The corresponding list of candidate regions in the XRQv2, PCS8 and LR1 genomes was identified by aligning 100bp flanking sequences around significant SNPs or k-mers from the source genome to the other reference genomes. Candidate genes within the defined regions were further explored for enrichment of gene ontology (GO) terms based on the information available in the annotation files (GFF). Enrichment analysis was performed using the org.At.tair.db (Carlson 2019) and ClusterProfiler v4.3.1 R packages (Wu et al., 2021) with p-value < 0.05 and q-value < 0.2. Significance level was adjusted to account for multiple testing using FDR < 0.05. To explore whether significant *k*-mers coincide within introgressions from wild species, we used the introgression maps available for the HA412HOv2, XRQv2, PCS8, and LR1 reference genomes (Huang et al., 2023).

### RNA-Seq data analysis

Raw reads from an RNA-seq experiment conducted for infested and non-infested roots of a resistant sunflower cultivar ‘EMEK3’ were obtained from a previous study (Sisou et al., 2021). Raw reads were trimmed and low-quality reads were filtered using Trimmomatic v.036 (Bolger et al., 2014), and inspected for quality using FastQC v.0.11.9 (Andrews, 2020). Cleaned high-quality reads were aligned to the HA412HOv2 genome using STAR v.2.5.2b (Dobin et al., 2013) and the expression levels were estimated and normalized to the number of reads per kilobase per million reads mapped (RPKM) using RSEM v.1.2.31 (Li & Dewey, 2011). Differential expression analysis was conducted with DESeq2 (Love et al., 2014) between infested and non-infested samples. Differentially expressed genes were considered as significant after correction for multiple testing at FDR < 0.05.

## Results

### High-throughput screening of broomrape resistance in the SAM population

Two *Orobanche cumana* populations were collected from infected sunflower fields in Yavor Farm (‘Yavor’) and Kibbutz Gadot (‘Gadot’) located 50 Km apart in northern Israel. Prior to use, seed viability was evaluated with 10^−5^M of GR24 and DCL to ensure a proper germination. Germination rates were higher with DCL (‘Yavor’ = 75.2%, ‘Gadot’ = 66.7%) than with GR24 (‘Yavor’ = 68.2%, ‘Gadot’ = 57.2%) and low spontaneous germination (‘Yavor’ = 1.7%, ‘Gadot’ = 1.2%) was observed under sterilized water (Fig. S1). To phenotype the root system of each accession in the SAM population, a high throughput rhizotron growth platform was developed, and an image analysis toolkit was employed (Fig. S2). A total of 1413 plants were screened in this system, comprising 251 and 220 accessions for the ‘Yavor’ and ‘Gadot’ *O. cumana* populations respectively, each with three independent replicates. The image analysis toolkit enabled to screen for different traits including number of attachments, area of each tubercle, classification of tubercles as healthy or necrotic, broomrape germination rate and sunflower root area. The most consistent phenotypes across replicates were the healthy attachment density (HealAD) which indicates an unsuccessful infection (or pre-haustorial resistance), and the necrotic attachments density (NecAD) which indicates a post-haustorial resistance to broomrape.

The average number of attachments of the ‘Yavor’ broomrape per sunflower plant was 43.05 of which 74% were healthy attachments and 26% necrotic attachments. For the ‘Gadot’ broomrape, the average number of attachments was 18.3 of which 47% were healthy attachments and 53% necrotic attachments. Altogether 66 and 95 accessions showed an absolute resistance to ‘Yavor’ and ‘Gadot’ broomrapes respectively, of which 34 accessions showed complete resistance to both populations (Table S1). Within the SAM population, a significant difference in the response to broomrape was observed between the four main genetic groups which distinguish maintainer and restorer breeding lines (HA and RHA), and between oil and confectionary varieties (HA-Oil, HA-NonOil, RHA-Oil, RHA-Non-Oil). Overall, the oil varieties were more resistant to both broomrape populations with marginal differences between HA-Oil and RHA-Oil for healthy or necrotic attachments (Fig. 1a, Table S2).

**Fig 1.**
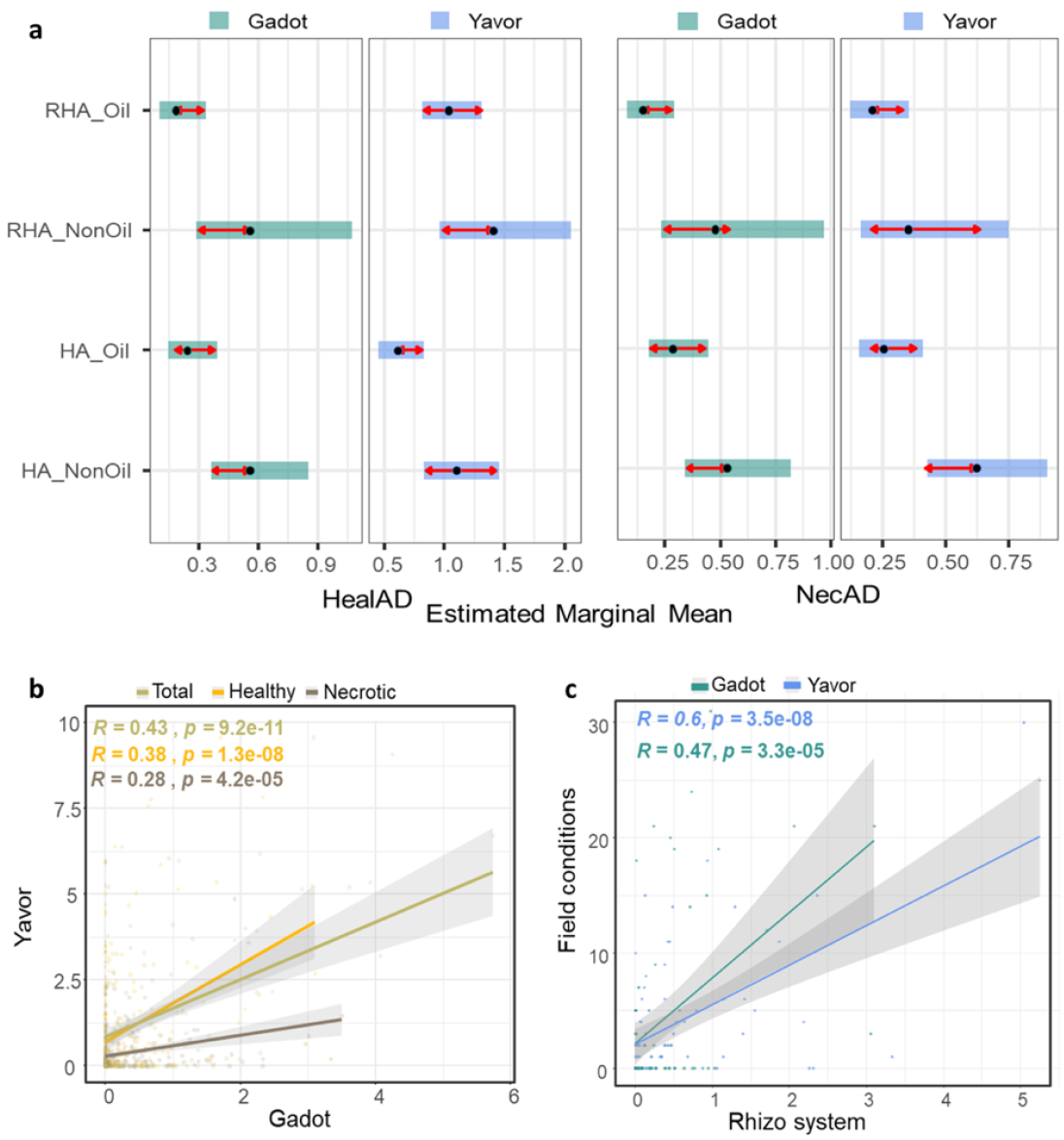
Evaluating resistance to broomrape races in the SAM population. (**a**) Comparisons of estimated marginal means (EMM) for HealAD and NecAD. Significant difference between sunflower breeding type is indicated with non-overlapping red arrows. (**b**) Pearson correlation tests between ‘Yavor’ and ‘Gadot’ races, and (**c**) Pearson correlation tests between the field (number of shoots) and in the rhizotron system (number of attachments) for both broomrape races.

The two broomrape populations were previously characterized as distinct races and evaluations conducted with a testing sunflower panel resulted with an unclear race definition for the Israeli populations due to high diversity in the response within each population. To evaluate the similarity between populations, a Pearson correlation was performed indicating a weak, yet significant correlation between the ‘Yavor’ and ‘Gadot’ broomrapes (Fig 1b). However, substantial and significant differences were observed between broomrape populations for the total number of attachments (*F* = 12.88, *p* = 3.81×10^−4^) and healthy attachments (*F* = 22.00, *p* = 3.97×10^−6^) but not for number of necrotic attachments (Table S3). Thus, overall, our results support previous virulence observations indicating the two populations can be defined as distinguished races with some overlap between them.

The rhizotron system was highly efficient for screening the large SAM population in a reasonable time frame, yet it is unclear how this system is predictive for the performance of fully grown sunflower plants under field conditions. To address this, we conducted two field experiments, one with each broomrape race, and phenotyped the plants performance under infection and control conditions. To obtain a manageable experimental setup, we selected from the SAM populations 75 accessions that were most susceptible and resistant to broomrape infection in the rhizotron screening of each broomrape race. Thus, a total of 1232 plots were grown for both ‘Yavor’ and ‘Gadot’ in the field experiments (616 per population). To evaluate the level of infection, the number of broomrape shoots was counted in each plot. Significant differences were observed in both fields for the measured traits and level of infection (Tables S4-S5), where stem diameter (*F*_*’Yavor’*_ = 38.93, *p* = 1.06×10^−9^ ; *F*_*’Gadot’*_ = 5.40, *p* = 0.02), and days to anthesis (*F*_*’Yavor’*_ = 2.63, *p* = 0.09; *F*_*’Gadot’*_ = 3.23, *p* = 0.07) showed the largest difference between infected and non-infected control plots (Table S5). To evaluate the consistency between the rhizotron system and field experiments, a Pearson correlation test was conducted for the number of attachments per plant (rhizotron) and the number of broomrape shoots counted (field) indicating a positive correlation for both ‘Yavor’ (*r*^*2*^ = 0.6, *p* = 3.50×10^−8^) and ‘Gadot’ (*r*^*2*^ = 0.47, *p* = 3.30×10^−5^) races, thus supporting the reliability of the rhizotron screening system (Fig. 1c).

### Resistance to broomrape is conferred by introgressions

To identify genomic regions associated with sunflower response to broomrape infection, genome-wide association analysis was conducted using the screening data from the rhizotron system for each broomrape race separately. A total of 251 and 220 accessions were screened for the ‘Yavor’ and ‘Gadot’ broomrapes, respectively. Inconsistency between replicates of specific accessions which showed no infestation in some plates and intensive infestation in others, has biased the distribution of phenotypes across accessions. To overcome this, we took an extreme phenotype mapping approach by extracting accessions at the edges of the distribution of each phenotype and converting their values to binary scores (Fig S3). Population structure and kinship matrices were calculated for each dataset indicating a consistent divergence between the four main breeding groups (Fig. S5). We used a univariate mixed linear model for each trait in each dataset (‘Yavor’/’Gadot’) separately. Significant associations were detected mainly for HealAD for both ‘Yavor and ‘Gadot’ (chromosomes 7, 9, 11, 13, 15, 16; Fig 2a) with very few overlapping signals between races, thus indicating a distinct genetic response to each broomrape race. Notably, only marginally significant associations were detected for NecAD in both broomrape races (Fig 2b).

**Fig 2.**
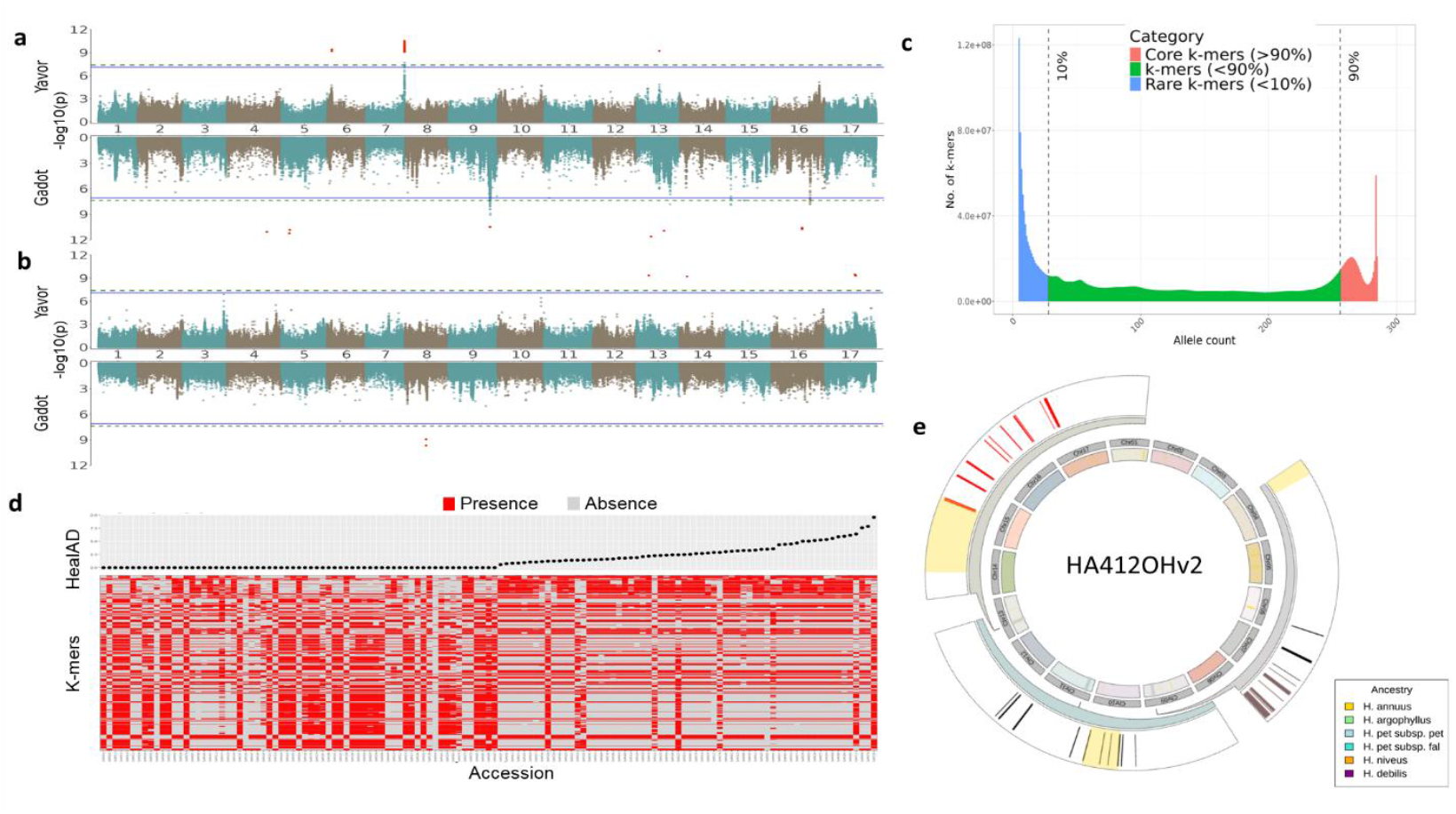
Identifying genomic regions that are associated with broomrape resistance. (**a**) Miami plots for maximum HealAD, and (**b**) NecAD in response to ‘Yavor’ (top) and ‘Gadot’ (bottom) races. Significantly associated *k*-mers are highlighted in red. Horizontal lines correspond to SimpleM (solid line) and Bonferroni (dashed line) correction for multiple testing at 5%. (**c**) Abundance distribution of *k*-mers in the SAM population. (**d**) Patterns of presence (red) absence (grey) variation of significantly associated *k*-mers (bottom) with HealAD phenotyps (top) measured for the ‘Yavor’ race. (**e**) Genomic regions that are significantly associated with broomrape resistance and overlap with introgressed region from wild relatives (outer circle) into the HA412OHv2 genome. Genomic regions identified with SNPs are colored in black and *k*-mers associations in red.

Despite their power in identifying genomic regions that are associated with a trait of interest, GWAS are limited to genetic variation that is detectable based on sequences in the reference genome. Previous work in sunflower and other species have demonstrated that a considerable fraction of genes is not represented in a single reference genome, specifically of introgressions that are associated with resistance to stress (Hübner et al., 2019). To expand our analyses, we screened sequencing reads from the entire SAM population to detect presence/absence variation (PAV) of *k*-mers (*k* = 31bp) that are associated with broomrape resistance. A total of 2,661,913,701 unique *k*-mers were identified across the SAM population, of which 21,095,787 *k*-mers were kept after filtering. Among the identified *k*-mers, 15,242,055 represent the core fraction that occurs across more than 90% of accessions, 22,482,716 *k*-mers were found in less than 10% of accessions (unique), and the remaining *k*-mers represent the dispensable fraction (Fig. 2c).

Next, the *k*-mers PAV information was converted into a genotype matrix and used for GWAS following similar procedure to the SNP-based GWAS. Altogether 660 *k*-mers were significantly associated with HealAD (‘Yavor’ = 554, ‘Gadot’ = 106) and 122 with NecAD (‘Yavor’ = 83, ‘Gadot’ = 39). To validate these results, the profile of *k*-mers PAV was explored across accessions, indicating a clear shift in *k*-mers PAV between susceptible and resistant accessions in the SAM population (Fig 2d, Fig S6). Identification of *k*-mers across accessions does not rely on a reference genome, hence their location along chromosomes is unknown. To further explore the identified associations, the position of the *k*-mers was determined by mapping them to the HA412OHv2 reference genome. To robustly identify the exact position of each *k*-mer, we used stringent alignment parameters with the cost of reducing the rate of *k*-mers mapped to the genome. To partially overcome this limitation without compromising for the stringent mapping, *k*-mers were also mapped to other available reference genomes (XRQv2, PSC8, LR1) which has increased the number of mapped *k*-mers and enabled to expand further our analyses. Among the 782 significant *k*-mers identified across all GWAS analyses, 168 were successfully mapped to the HA412OHv2 reference genome, 185 *k*-mers were mapped to XRQv2, 357 to the PSC8 genome, and 180 to the LR1 genome. Surprisingly, only 30 *k*-mers were successfully mapped to all genomes which implies that the significantly associated *k*-mers with response to broomrape infection is attributed to the dispensable fraction of the sunflower pangenome (Fig. S7).

Next, we explored overlaps between the coordinates of significant *k*-mers on the HA412OHv2 genome and significant signals obtained from the SNPs-based GWAS (Fig. 2a,b). Strong signals of association with HealAD, supported by SNPs and *k*-mers were observed on chromosome 7 and 9 for the ‘Yavor’ and ‘Gadot’ respectively, which further supports their definition as distinct races. Other associations were identified on chromosomes 6 in response to the ‘Yavor’ race, on chromosomes 4, 5, and 16 in response to ‘Gadot’, and a shared signal for both races on chromosome 13. No significant SNPs were identified for the NecAD trait, yet several significant *k*-mers were identified on chromosomes 8 (‘Gadot’), 13, 14 and 17 (‘Yavor’) (Fig. 2a,b, Table S6). These results demonstrate the power of non-reference *k*-mers in identifying genetic variation of potential importance.

To further explore whether the identified *k*-mers associations correspond to introgressions into cultivated sunflower, we compared the coordinates of significant *k*-mers to available maps of introgressions from six wild relative species (*H. annuus, H. argophyllus, H. petiolaris* subsp. *petiolaris, H. petiolaris* subsp. *fallax, H. niveus* and *H. debilis*). Among *k*-mers, 249 were mapped to genomic regions that coincide introgressions from wild relatives across the chromosomes of the inspected reference genomes (Fig 2e, Fig S8). Overall, we identified 16 regions that are associated with broomrape resistance and were introgressed from wild *H. annuus*, one from *H. debilis* and two from *H. petiolaris* (subsp. *petiolaris* and *fallax*).

### Introgressions of resistance genes from wild sunflowers

To further explore candidate genes that regulate the response of sunflower to broomrape infestation and their functional annotation, we extracted genes within 250Kbp up/down to significant signals (SNPs and *k*-mers) based on the HA412OHv2 genome annotation. This distance concord with the expected linkage disequilibrium decay in cultivated sunflower (Fig. S9). A total of 474 genes were detected, of which 386 genes are associated with HealAD (112 for ‘Yavor’ and for 274 ‘Gadot’), and 88 with NecAD (79 for ‘Yavor’ and 9 for ‘Gadot’) (Table S6). To illuminate the main biological processes involved in the sunflower response to broomrape infestation, a gene ontology (GO) enrichment and pathway analyses were conducted. As expected, the main GO terms that were significantly enriched include activation and regulation of the immune system, regulation of defense response, induced systemic resistance, symbiont-induced programmed cell death, response to biotic stimulus, phenylpropanoid and metabolic process, jasmonic acid mediated signaling pathway, and kinase activities (Fig. 3a, Fig. S10, Table S7). Among the identified genes, 55 are associated with pathogen-triggered innate immunity responses of which 17 are kinase RLK-Pelle genes, 2 serine/threonine protein kinases, 4 wall-associated receptor kinase, 2 enhanced disease resistance proteins, 7 Putative leucine-rich repeat-containing proteins, and 31 transcription factors of known function in biotic stress (MYB, C2H2, bZIP, and bHLH).

**Fig 3.**
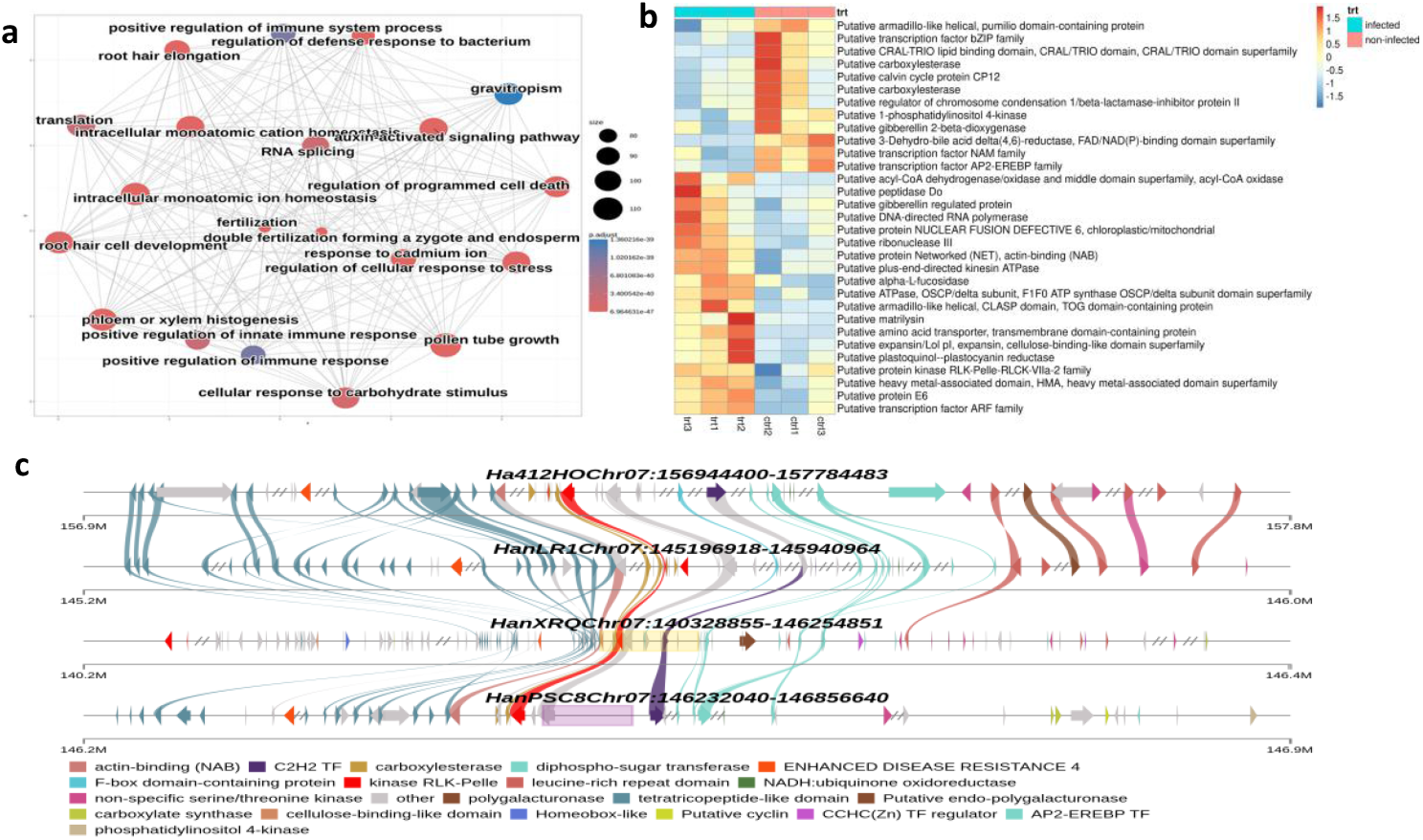
Exploration of genes that regulate the response of sunflower to broomrape. (**a**) GO enrichment pathway map of annotated genes associated with NecAD in response to ‘Yavor’ race; (**b**) Differentially expressed genes that were identified in the GWAS and RNA-Seq analyses. The top bar indicate the broomrape infestation (treatment) versus control, and the heatmap colors are from blue (down regulation) to red (up regulation). (**c**) clusters of genes mapped to the different reference genomes on chromosome 7 which contain numerous resistant genes nearby introgressed regions from wild *H. annuus* (marked in yellow bar).

Moreover, at least 17 SNPs and *k*-mers were located within coding sequences of genes including in key transcription factors (VOZ, C2H2). Substantial portion of the identified candidate genes were clustered in two genomic regions: one located on chromosome 7 (Ha412HOChr07:156944400-157784483) which is mainly associated in the ‘Yavor’ race, and another cluster on chromosome 9 (Ha412HOChr09:167075448-168109504) that is associated with the response in the ‘Gadot’ race (Fig S13). To expand the representation of genetic variation, we mapped these regions to all other reference genomes. Interestingly, the cluster on chromosome 7 was identified based on the XRQv2 reference as an introgression from wild *H. annuus* (HanXRQChr07:140328855-146254851) which harbors at least 7 resistance genes tightly linked to each other. Moreover, the same region in the PSC8 genome highlighted an introgression from *H. debilis* (HanPSC8Chr07:146232040-146856640) that contains a putative protein kinase RLK-Pelle-CrRLK1L-1, a carboxylesterase, and C2H2 transcription factor (Fig. 3c). Similarly, the genes cluster on chromosome 9 was identified in the LR1 (LR1Chr09:157163610-157985684) and XRQv2 (XRQv2Chr09:158290146-160953558) reference genomes as an introgression from *H. annuus* which contains at least 8 resistance genes including a putative protein kinase RLK-Lelle-LRR-Xa gene, an enhanced disease resistance gene (EDR2), AP2-EREBP transcription factor and others (Table S8, Fig. S13).

To further evaluate the functional response of the identified candidate genes, we analyzed RNASeq data available for the ‘EMEK3’, a resistant variety to the ‘Yavor’ race (Sisou et al., 2021). Altogether, 4731 significant differentially expressed genes were identified (Fig. S12, Table S9) of which 38 were also identified in the GWAS analyses (Fig 3b). Among these genes, several were mapped to the gene cluster on chromosome 9 including the putative protein kinase and putative protein expansin, thus indicating a functional activity of genes that were introgressed from wild species into the cultivated sunflower genome (Fig. 3b, Fig. S13).

## Discussion

Since the early stages of domestication, farmers have been troubled by yield loss due to pathogens infestation, thus sparkling a persistent and ongoing arms race (Lebeda & Burdon, 2023; Stukenbrock & McDonald, 2008). Parasitic plants pose a specific challenge due to their biological similarity to the host which impedes the use of many chemical and agrotechnical practices (Eizenberg et al., 2009; 2014). Therefore, breeding towards resistance to parasitic plants is the most effective and environmentally friendly approach to overcome the severe damage imposed by broomrape infestation. In this study, we explored the genetic response of sunflower to the broomrape parasitic plant (*Orobanche cummana*) which is alarmingly spreading across continents, leading to significant yield losses (Hu et al., 2020). So far, several QTLs that are associated with resistance to broomrape have been identified, yet the exact genes conferring resistance have been discovered only in the case of *Or7* (Duriez et al., 2019). Most studies, have used a bi-parental segregating populations and a limited set of genetic markers, thus limiting the discovery of resistance genes at fine resolution (Bergelson & Roux, 2010; Zhu et al., 2008). To overcome these limitations, we have developed a high-resolution rhizotron screening platform for the root system (Fig S2) supplemented with an image analysis phenotyping approach (Hasson et al. 2021). This approach enables to screen large diversity panels efficiently and to implement large GWAS for exploring resistance to parasitic plants in crop species. This platform facilitated the analysis of the sunflower association mapping (SAM) population which has been intensively genotyped and explored for biotic and abiotic stress resistance (Gao et al. 2019; Hübner et al. 2019; Mandel et al. 2013; Todesco et al. 2020). Despite the rhizotron platform efficiency, it has some limitations due to variation in the germination and growth rates between accessions, the limited space for root growth in the plate, and the sensitivity of broomrape to the hydroponic conditions. Nevertheless, we were able to obtain high quality data to perform GWAS for broomrape resistance across a wide diversity panel. Comparing the rhizotron phenotypes to field trials indicated that the system largely predicts the response of sunflower to broomrape infection in an agricultural field setup (Fig 1d). Due to the challenges of the rhizotron system, we focused on two traits for which consistent results were obtained across replicates: the number of healthy and necrotic tubercles which represent susceptibility (pre-attachment) and post-haustorial resistance, respectively (Fig. 1). These traits were further used in the GWAS and indicated a different genetic response. Overall, a stronger association signal was obtained for the healthy tubercles in all analyses, presumably because post-haustorial resistance is less common in sunflower (Martín-Sanz et al., 2020).

Based on the comparison between the response to the ‘Yavor’ and ‘Gadot’ populations and the differences in the rhizotron and fields setups we support previous observations indicating that these populations are assigned to different races (Eizenberg et al., 2004). The current racial nomenclature of broomrape (named A-H) follows the response of sunflower differential lines to infestation. However, accumulating evidence support the geographical context of race development, thus suggesting a different classification approach (Calderón-González et al., 2024). Our results largely support the geographical context of racial definition for broomrape populations, as new races tend to evolve locally from existing races (Dor et al., 2020; Joel et al. 2011).

The GWAS was implemented for each race separately and highlighted a signal on chromosome 7, around the Or7 QTL (Duriez et al., 2019) for the ‘Yavor’ race, and on chromosome 9, 15 and 16 for the ‘Gadot’ race. These signals colocalize with QTLs identified previously inresponse to infestation with races E, F or G using bi-parent populations (Imerovski et al., 2019; Louarn et al., 2016). Despite the higher mapping resolution obtained with the SAM population, the variants were called against a single reference genome (HA412), thus some genetic variation that underlies the response to broomrape infection in not represented (Haung et al. 2023; Hübner et al., 2019; Hübner 2022a). To overcome this, we implemented a *k*-mer based GWAS approach which enabled to detect additional signals on chromosomes 4, 6, 8, 13, 14 and 17 (Fig. 2). The *k*-mer GWAS approach is specifically relevant for the identification of introgressions that are not necessarily represented in a single reference genome and were brought from wild relatives to confer pathogen resistance (Huang et al., 2023; Hübner et al., 2019). Indeed, 20 of the significantly associated *k*-mers were found in genomic regions that were introgressed into cultivated sunflower from wild *Helianthus* species. Thus, emphasizing the importance of expanding the genomic mapping analyses beyond standard SNP-based GWAS. For example, the genomic region identified on chromosome 7 is close to the *Or7* locus which was mapped using a bi-parental population and confers resistance to the F broomrape race (Duriez et al., 2019). Exploring this genomic region in other reference genomes enabled to expand the representation of genetic variation highlighting a cluster of resistance genes, including introgressions from *H. annuus* and *H. debilis* (Fig S13; Table S8). Among the identified introgressions from *H. debilis* we detected a putative protein kinase RLK-Pelle-CrRLK1L-1, a carboxylesterase and C2H2 transcription factor, thus suggesting that resistance may be conferred also by other genes within the same cluster in addition to the identified leucine-rich repeat receptor-like kinase (Duriez et al., 2019). We also found multiple sugar transferase and glycosyl transferase were recently described as important factors in conferring resistance to broomrape in sunflower (Calderón-González et al., 2023).

Broomrape is quickly spreading across fields despite its high sensitivity and reliance on a very specific chemical recognition of the host. Interestingly, the pathogenic broomrape evolved outside the core region of wild sunflower and related *Helianthus* species, and specific recognition mechanism of wild sunflowers is hindered. Therefore, genetic variation from wild relatives is highly relevant to confer broomrape resistance in cultivated sunflower. Interestingly, a recent evaluation of invasive wild sunflower in Israel (Hübner et al. 2022b) also highlighted resistance mechanism specifically to the Israeli broomrape races (data not shown).

This research presents a novel high-throughput platform for screening large diversity panels, such as the SAM population. Our approach successfully identified key genetic variations and specific sunflower accessions with resistance to various broomrape races. The availability of resistance genes sourced from wild sunflower relatives is highly promising for breeding efforts. However, the potential spread of broomrape to new regions, like North America, may accelerate the evolution of the parasite, challenging the effectiveness of these wild resistance genes. This ongoing co-evolutionary “arms race” highlights the importance of continually incorporating genetic diversity from wild relatives to maintain a breeding advantage.

## Acknowledgements

We thank Guy Atzmon, Amit Wallach and Hod Hasson for their help with preparation of the field experiment and phenotyping. We also thank ‘Sha’ar Ha’amakim Seeds Ltd.’ for providing susceptible and resistant seeds used as control. This research was supported by a grant number 21-01-0034 (SH and HE) from the Ministry of Agriculture, Israel.

## Author contributions

HE and SH conceived and supervised the research project. DS planned and designed the experiments and performed the data analyses. HZ, MEW and DS performed the experiments and measured phenotypes. DS and SH wrote the first draft, and all authors revised and contributed to the final version of the manuscript.

## Supporting Information

Fig. S1. Broomrape seed germination evaluation

Fig. S2. Phenotyping platform for sunflower broomrape resistance

Fig. S3. Phenotype distribution and thresholds for the extreme phenotype approach

Fig. S4. Field evaluation

Fig. S5. Population structure and kinship matrices of the SAM population

Fig. S6. PAV patterns of significant k-mers

Fig. S7. Shared k-mers among HA412OH, XRQ, PSC8 and LR1 genomes

Fig. S8 Introgressions of genes associated with broomrape resistance from different wild relatives in different referenced genomes

Fig. S9. LD-decay of the SAM population

Fig. S10. GO enrichment analysis of genes associated with broomrape resistance

Fig. S11. Volcano plot of DEGs for broomrape resistance

Fig. S12. Ven diagram showing shared genes among analyses

Fig. S13. Comparison of genomic regions clusters associated with broomrape resistance across reference genomes

Table S1. Summary of SAM population response to sunflower broomrape

Table S2. Summary table for the analysis of variance between the four main genetic groups among the SAM population in response to Yavor and Gadot broomrapes

Table S3. Summary table for the analysis of variance between Yavor and Gadot broomrapes Table S4. Summary table for the analysis of variance of the field experiment

Table S5. Summary of phenotypic data collected in the field experiment for Yavor and Gadot broomrape populations

Table S6. SNPs and k-mers significantly associated with broomrape resistance and annotated genes in HA412OH, XRQ, PSC8 and LR1 genomes

Table S7. Significantly enriched GO terms of annotated genes associated with response to broomrape infection

Table S8. Annotated genes of significantly associated markers coincide Introgressed regions in HA412OH, XRQ, PSC8 and LR1 genomes

Table S8. Differentially expressed genes of infested and non-infested roots of a resistant sunflower cultivar ‘EMEK3’

*Tables S5 – S8 Provided as an Table_data.xlsx document

## Notes

### Competing Interest Statement

The authors have declared no competing interest.

## References

Akhtouch, B., del Moral, L., Leon, A., Velasco, L., Fernández-Martínez, J. M., & Pérez-Vich, B. (2016). Genetic study of recessive broomrape resistance in sunflower. Euphytica, 209, 419–428. 10.1007/s10681-016-1652-z

Amanat, S., Requena, T., & Lopez-Escamez, J. A. (2020). A systematic review of extreme phenotype strategies to search for rare variants in genetic studies of complex disorders. In Genes (Vol. 11, Issue 9). 10.3390/genes11090987

Andrews, S. (2020). Babraham Bioinformatics - FastQC A Quality Control tool for High Throughput Sequence Data. https://github.com/s-andrews/FastQC.

Bates, D., Mächler, M., Bolker, B. M., & Walker, S. C. (2015). Fitting linear mixed-effects models using lme4. Journal of Statistical Software, 67(1). 10.18637/jss.v067.i01

Bergelson, J., & Roux, F. (2010). Towards identifying genes underlying ecologically relevant traits in Arabidopsis thaliana. In Nature Reviews Genetics (Vol. 11, Issue 12). 10.1038/nrg2896

Bolger, a. M., Lohse, M., & Usadel, B. (2014). Trimmomatic: A flexible read trimming tool for Illumina NGS data. Bioinformatics, 30(15).

Calderón-González, Á., Fernández-Melero, B., del Moral, L., Muños, S., Velasco, L., & Pérez-Vich, B. (2024). Mapping an avirulence gene in the sunflower parasitic weed Orobanche cumana and characterization of host selection based on virulence alleles. BMC Plant Biology, 24. 10.1186/s12870-024-05855-2

Calderón-González, Á., Pérez-Vich, B., Pouilly, N., Boniface, M. C., Louarn, J., Velasco, L., & Muños, S. (2023). Association mapping for broomrape resistance in sunflower. Frontiers in Plant Science, 13. 10.3389/fpls.2022.1056231

Carlson Marc. (2019). GO.db: A Set of Annotation Maps Describing the Entire Gene Ontology (. R package version 3.10.10). 10.18129/B9.bioc.GO.db

Chang, C. C., Chow, C. C., Tellier, L. C. A. M., Vattikuti, S., Purcell, S. M., & Lee, J. J. (2015). Second-generation PLINK: Rising to the challenge of larger and richer datasets. GigaScience, 4(1). 10.1186/s13742-015-0047-8

Cvejić, S., Radanović, A., Dedić, B., Jocković, M., Jocić, S., & Miladinović, D. (2020). Genetic and genomic tools in sunflower breeding for broomrape resistance. In Genes (Vol. 11). MDPI AG. 10.3390/genes11020152

Danecek, P., Bonfield, J. K., Liddle, J., Marshall, J., Ohan, V., Pollard, M. O., Whitwham, A., Keane, T., McCarthy, S. A., & Davies, R. M. (2021). Twelve years of SAMtools and BCFtools. GigaScience, 10(2). 10.1093/gigascience/giab008

De Zélicourt, A., Letousey, P., Thoiron, S., Campion, C., Simoneau, P., Elmorjani, K., Marion, D., Simier, P., & Delavault, P. (2007). Ha-DEF1, a sunflower defensin, induces cell death in Orobanche parasitic plants. Planta, 226, 591–600. 10.1007/s00425-007-0507-1

Demirjian, C., Vailleau, F., Berthomé, R., & Roux, F. (2023). Genome-wide association studies in plant pathosystems: success or failure? In Trends in Plant Science (Vol. 28, Issue 4). 10.1016/j.tplants.2022.11.006

Dobin, A., Davis, C. A., Schlesinger, F., Drenkow, J., Zaleski, C., Jha, S., Batut, P., Chaisson, M., & Gingeras, T. R. (2013). STAR: Ultrafast universal RNA-seq aligner. Bioinformatics, 29(1). 10.1093/bioinformatics/bts635

Duca, M., Clapco, S., & Joita-Pacureanu, M. (2022). Racial status of Orobanche cumana Wallr. in some countries other the world. In Helia (Vol. 45, pp. 1–22). De Gruyter Open Ltd. 10.1515/helia-2022-0002

Duriez, P., Vautrin, S., Auriac, M. C., Bazerque, J., Boniface, M. C., Callot, C., Carrère, S., Cauet, S., Chabaud, M., Gentou, F., Lopez-Sendon, M., Paris, C., Pegot-Espagnet, P., Rousseaux, J. C., Pérez-Vich, B., Velasco, L., Bergès, H., Piquemal, J., & Muños, S. (2019). A receptor-like kinase enhances sunflower resistance to Orobanche cumana. Nature Plants, 5, 1211–1215. 10.1038/s41477-019-0556-z

Echevarría-Zomeño, S., Pérez-De-Luque, A., Jorrín, J., & Maldonado, A. M. (2006). Pre-haustorial resistance to broomrape (Orobanche cumana) in sunflower (Helianthus annuus): Cytochemical studies. Journal of Experimental Botany, 57, 4189–4200. 10.1093/jxb/erl195

Eizenberg, H., Hershenhorn, J., & Ephrath, J. E. (2009). Factors affecting the efficacy of Orobanche cumana chemical control in sunflower. Weed Research, 49, 308–315. 10.1111/j.1365-3180.2009.00701.x

Eizenberg, H., Hershenhorn, J., Ephrath, J. H., & Kanampiu, F. (2014). Chemical control. In Parasitic Orobanchaceae: Parasitic Mechanisms and Control Strategies. Springer-Verlag Berlin Heidelberg. 10.1007/978-3-642-38146-1_23

Eizenberg, H., Plakhine, D., Landa, T., Achdari, G., Joel, D. M., & Hershenhorn, J. (2004). First Report of a New Race of Sunflower Broomrape (Orobanche cumana) in Israel. Plant Disease, 88(11). 10.1094/pdis.2004.88.11.1284c

Fernández-Aparicio, M., del Moral, L., Muños, S., Velasco, L., & Pérez-Vich, B. (2022). Genetic and physiological characterization of sunflower resistance provided by the wild-derived OrDeb2 gene against highly virulent races of Orobanche cumana Wallr. Theoretical and Applied Genetics, 135, 501–525. 10.1007/s00122-021-03979-9

Fernández-Martínez, J. M., Pérez-Vich, B., & Velasco, L. (2015). Sunflower Broomrape (Orobanche cumana Wallr.). In Sunflower: Chemistry, Production, Processing, and Utilization (pp. 129–155). Elsevier Inc. 10.1016/B978-1-893997-94-3.50011-8

Fernández-Melero, B., del Moral, L., Todesco, M., Rieseberg, L. H., Owens, G. L., Carrère, S., Chabaud, M., Muños, S., Velasco, L., & Pérez-Vich, B. (2024). Development and characterization of a new sunflower source of resistance to race G of Orobanche cumana Wallr. derived from Helianthus anomalus. Theoretical and Applied Genetics, 137. 10.1007/s00122-024-04558-4

Fox, J., & Weisberg, S. (2019). An {R} Companion to Applied Regression, Third Edition. Thousand Oaks CA: Sage., September 2012.

Gao, X. (2011). Multiple testing corrections for imputed SNPs. Genetic Epidemiology, 35, 154–158. 10.1002/gepi.20563

Goldwasser, Y., Kleifeld, Y., Golan, S., Bargutti, A., & Rubin, B. (1995). Dissipation of metham‐sodium from soil and its effect on the control of Orobanche aegyptiaca. Weed Research, 35(6). 10.1111/j.1365-3180.1995.tb01641.x

Hoagland D.R, & Arnon D.I. (1950). The water-culture method for growing plants without soil. Univ. of California, College of Agriculture, 1938, 347, No. 2nd edit, 32.

Hothorn, T., Bretz, F., & Westfall, P. (2008). Simultaneous inference in general parametric models. In Biometrical Journal (Vol. 50, Issue 3). 10.1002/bimj.200810425

Hu, L., Wang, J., Yang, C., Islam, F., Bouwmeester, H. J., Muños, S., & Zhou, W. (2020). The effect of virulence and resistance mechanisms on the interactions between parasitic plants and their hosts. International Journal of Molecular Sciences, 21(23). 10.3390/ijms21239013

Huang, K., Jahani, M., Gouzy, J., Legendre, A., Carrere, S., Lázaro-Guevara, J. M., González Segovia, E. G., Todesco, M., Mayjonade, B., Rodde, N., Cauet, S., Dufau, I., Staton, S. E., Pouilly, N., Boniface, M. C., Tapy, C., Mangin, B., Duhnen, A., Gautier, V., … Rieseberg, L. H. (2023). The genomics of linkage drag in inbred lines of sunflower. Proceedings of the National Academy of Sciences of the United States of America, 120(14). 10.1073/pnas.2205783119

Hübner, S., Bercovich, N., Todesco, M., Mandel, J. R., Odenheimer, J., Ziegler, E., Lee, J. S., Baute, G. J., Owens, G. L., Grassa, C. J., Ebert, D. P., Ostevik, K. L., Moyers, B. T., Yakimowski, S., Masalia, R. R., Gao, L., Ćalić, I., Bowers, J. E., Kane, N. C., … Rieseberg, L. H. (2019). Sunflower pan-genome analysis shows that hybridization altered gene content and disease resistance. Nature Plants, 5, 54–62. 10.1038/s41477-018-0329-0

Imerovski, I., Dedić, B., Cvejić, S., Miladinović, D., Jocić, S., Owens, G. L., Tubić, N. K., & Rieseberg, L. H. (2019). BSA-seq mapping reveals major QTL for broomrape resistance in four sunflower lines. Molecular Breeding, 39. 10.1007/s11032-019-0948-9

Imerovski, I., Dimitrijević, A., Miladinović, D., Dedić, B., Jocić, S., Tubić, N. K., & Cvejić, S. (2016). Mapping of a new gene for resistance to broomrape races higher than F. Euphytica, 209, 281– 289. 10.1007/s10681-015-1597-7

Joel, D. M., Chaudhuri, S. K., Plakhine, D., Ziadna, H., & Steffens, J. C. (2011). Dehydrocostus lactone is exuded from sunflower roots and stimulates germination of the root parasite Orobanche cumana. Phytochemistry, 72, 624–634. 10.1016/j.phytochem.2011.01.037

Kokot, M., Dlugosz, M., & Deorowicz, S. (2017). KMC 3: counting and manipulating k-mer statistics. Bioinformatics (Oxford, England), 33(17). 10.1093/bioinformatics/btx304

Lebeda, A., & Burdon, J. J. (2023). Studying Wild Plant Pathosystems to Understand Crop Plant Pathosystems: Status, Gaps, Challenges, and Perspectives. Phytopathology, 113(3). 10.1094/PHYTO-01-22-0018-PER

Lenth, R. V. (2024). emmeans: Estimated Marginal Means, aka Least-Squares Means. R Package Version 1.10.2.090002.

Letousey, P., De Zélicourt, A., Vieira Dos Santos, C., Thoiron, S., Monteau, F., Simier, P., Thalouarn, P., & Delavault, P. (2007). Molecular analysis of resistance mechanisms to Orobanche cumana in sunflower. Plant Pathology, 56, 536–546. 10.1111/j.1365-3059.2007.01575.x

Li, B., & Dewey, C. N. (2011). RSEM: Accurate transcript quantification from RNA-Seq data with or without a reference genome. BMC Bioinformatics, 12. 10.1186/1471-2105-12-323

Li, H., & Durbin, R. (2009). Fast and accurate short read alignment with Burrows-Wheeler transform. Bioinformatics, 25(14). 10.1093/bioinformatics/btp324

Li, Y., Levran, O., Kim, J. J., Zhang, T., Chen, X., & Suo, C. (2019). Extreme sampling design in genetic association mapping of quantitative trait loci using balanced and unbalanced case-control samples. Scientific Reports, 9(1). 10.1038/s41598-019-51790-w

Louarn, J., Boniface, M. C., Pouilly, N., Velasco, L., Pérez-Vich, B., Vincourt, P., & Muños, S. (2016). Sunflower resistance to broomrape (Orobanche cumana) is controlled by specific qtls for different parasitism stages. Frontiers in Plant Science, 7. 10.3389/fpls.2016.00590

Love, M. I., Huber, W., & Anders, S. (2014). Moderated estimation of fold change and dispersion for RNA-seq data with DESeq2. Genome Biology, 15(12). 10.1186/s13059-014-0550-8

Lu, Y. H., Gagne, G., Grezes-Besset, B., & Blanchard, P. (1999). Integration of a molecular linkage group containing the broomrape resistance gene Or5 into an RFLP map in sunflower. Genome, 42(3). 10.1139/g98-135

Lu, Y. H., Melero-Vara, J. M., García-Tejada, J. A., & Blanchard, P. (2000). Development of SCAR markers linked to the gene Or5 conferring resistance to broomrape (Orobanche cumana Wallr.) in sunflower. Theoretical and Applied Genetics, 100(3–4). 10.1007/s001220050083

Mandel, J. R., Dechaine, J. M., Marek, L. F., & Burke, J. M. (2011). Genetic diversity and population structure in cultivated sunflower and a comparison to its wild progenitor, Helianthus annuus L. Theoretical and Applied Genetics, 123, 693–704. 10.1007/s00122-011-1619-3

Mandel, J. R., Nambeesan, S., Bowers, J. E., Marek, L. F., Ebert, D., Rieseberg, L. H., Knapp, S. J., & Burke, J. M. (2013). Association Mapping and the Genomic Consequences of Selection in Sunflower. PLoS Genetics, 9. 10.1371/journal.pgen.1003378

Martín-Sanz, A., Pérez-Vich, B., Rueda, S., Fernández-Martínez, J. M., & Velasco, L. (2020). Characterization of post-haustorial resistance to sunflower broomrape. Crop Science, 60, 1188– 1198. 10.1002/csc2.20002

Molinero-Ruiz, L., Delavault, P., Pérez-Vich, B., Pacureanu-Joita, M., Bulos, M., Altieri, E., & Domínguez, J. (2015). History of the race structure of Orobanche cumana and the breeding of sunflower for resistance to this parasitic weed: A review. In Spanish Journal of Agricultural Research (Vol. 13, Issue 4). 10.5424/sjar/2015134-8080

Parker, C. (2009). Observations on the current status of orobanche and striga problems worldwide. In Pest Management Science (Vol. 65, Issue 5). 10.1002/ps.1713

Pérez-de-Luque, A., González-Verdejo, C. I., Lozano, M. D., Dita, M. A., Cubero, J. I., González-Melendi, P., Risueño, M. C., & Rubiales, D. (2006). Protein cross-linking, peroxidase and β-1,3-endoglucanase involved in resistance of pea against Orobanche crenata. Journal of Experimental Botany, 57, 1461–1469. 10.1093/jxb/erj127

Pérez-De-Luque, A., Lozano, M. D., Moreno, M. T., Testillano, P. S., & Rubiales, D. (2007). Resistance to broomrape (Orobanche crenata) in faba bean (Vicia faba): Cell wall changes associated with prehaustorial defensive mechanisms. Annals of Applied Biology, 151, 89–98. 10.1111/j.1744-7348.2007.00164.x

Pérez-Vich, B., Akhtouch, B., Knapp, S. J., Leon, A. J., Velasco, L., Fernández-Martínez, J. M., & Berry, S. T. (2004). Quantitative trait loci for broomrape (Orobanche cumana Wallr.) resistance in sunflower. Theoretical and Applied Genetics, 109, 92–102. 10.1007/s00122-004-1599-7

Plakhine, D., & Joel, D. M. (2010). Ecophysiological consideration of orobanche cumana germination. Helia, 33(52). 10.2298/HEL1052013P

Pubert, C., Boniface, M. C., Legendre, A., Chabaud, M., Carrère, S., Callot, C., Cravero, C., Dufau, I., Patrascoiu, M., Baussart, A., Belmonte, E., Gautier, V., Poncet, C., Zhao, J., Hu, L., Zhou, W., Langlade, N., Vautrin, S., Coussy, C., & Muños, S. (2024). A cluster of putative resistance genes is associated with a dominant resistance to sunflower broomrape. Theoretical and Applied Genetics, 137. 10.1007/s00122-024-04594-0

Schneider, C. A., Rasband, W. S., & Eliceiri, K. W. (2012). NIH Image to ImageJ: 25 years of image analysis. In Nature Methods (Vol. 9, pp. 671–675). 10.1038/nmeth.2089

Seiler, G. J., & Jan, C. C. (2014). Wild sunflower species as a genetic resource for resistance to sunflower broomrape (orobanche cumana Wallr.). Helia, 37, 129–139. 10.1515/helia-2014-0013

Sisou, D., Tadmor, Y., Plakhine, D., Ziadna, H., Hübner, S., & Eizenberg, H. (2021). Biological and transcriptomic characterization of pre-haustorial resistance to sunflower broomrape (Orobanche cumana w.) in sunflowers (helianthus annuus). Plants, 10. 10.3390/plants10091810

Stukenbrock, E. H., & McDonald, B. A. (2008). The origins of plant pathogens in agro-ecosystems. In Annual Review of Phytopathology (Vol. 46). 10.1146/annurev.phyto.010708.154114

Tang, S., Heesacker, A., Kishore, V. K., Fernandez, A., Sadik, E. S., Cole, G., & Knapp, S. J. (2003). Genetic mapping of the Or5 gene for resistance to Orobanche Race E in sunflower. Crop Science, 43, 1021–1028. 10.2135/cropsci2003.1021

Todesco, M., Owens, G. L., Bercovich, N., Légaré, J. S., Soudi, S., Burge, D. O., Huang, K., Ostevik, K. L., Drummond, E. B. M., Imerovski, I., Lande, K., Pascual-Robles, M. A., Nanavati, M., Jahani, M., Cheung, W., Staton, S. E., Muños, S., Nielsen, R., Donovan, L. A., … Rieseberg, L. H. (2020). Massive haplotypes underlie ecotypic differentiation in sunflowers. Nature, 584, 602–607. 10.1038/s41586-020-2467-6

Voichek, Y., & Weigel, D. (2020). Identifying genetic variants underlying phenotypic variation in plants without complete genomes. Nature Genetics, 52, 534–540. 10.1038/s41588-020-0612-7

Vrânceanu A.V, Tudor V. A, Stoenescu F. M, & Pirvu N. (1980, June 8). Virulence groups of Orobanche cumana Wallr. differential hosts and resistance sources and genes in sunflower. https://Www.Cabidigitallibrary.Org/Doi/Full/10.5555/19831624372.

Wilkinson, L. (2011). ggplot2: Elegant Graphics for Data Analysis by WICKHAM, H. Biometrics, 67(2). 10.1111/j.1541-0420.2011.01616.x

Wu, T., Hu, E., Xu, S., Chen, M., Guo, P., Dai, Z., Feng, T., Zhou, L., Tang, W., Zhan, L., Fu, X., Liu, S., Bo, X., & Yu, G. (2021). clusterProfiler 4.0: A universal enrichment tool for interpreting omics data. Innovation, 2. 10.1016/j.xinn.2021.100141

Yoder, J. I., & Scholes, J. D. (2010). Host plant resistance to parasitic weeds; recent progress and bottlenecks. In Current Opinion in Plant Biology (Vol. 13, Issue 4). 10.1016/j.pbi.2010.04.011

Yoshida, S., & Shirasu, K. (2012). Plants that attack plants: Molecular elucidation of plant parasitism. In Current Opinion in Plant Biology (Vol. 15, Issue 6). 10.1016/j.pbi.2012.07.004

Zhang, C., Dong, S. S., Xu, J. Y., He, W. M., & Yang, T. L. (2019). PopLDdecay: A fast and effective tool for linkage disequilibrium decay analysis based on variant call format files. Bioinformatics, 35, 1786–1788. 10.1093/bioinformatics/bty875

Zhou, X., & Stephens, M. (2012). Genome-wide efficient mixed-model analysis for association studies. Nature Genetics, 44, 821–824. 10.1038/ng.2310

Zhu, C., Gore, M., Buckler, E. S., & Yu, J. (2008). Status and Prospects of Association Mapping in Plants. The Plant Genome, 1(1). 10.3835/plantgenome2008.02.0089

