## Supplemental_information for "Wild genes to the rescue: High-throughput genomics reveals the wild source of broomrape resistance in sunflower"

### **Supporting information**

Fig. S1. Broomrape seed germination evaluation

Fig. S2. Phenotyping platform for sunflower broomrape resistance

Fig. S3. Phenotype distribution and thresholds for the extreme phenotype approach

Fig. S4. Field evaluation

Fig. S5. Population structure and kinship matrices of the SAM population

Fig. S6. PAV patterns of significant k-mers

Fig. S7. Shared k-mers among HA412OH, XRQ, PSC8 and LR1 genomes

Fig. S8 Introgressions of genes associated with broomrape resistance from different wild relatives in different referenced genomes

Fig. S9. LD-decay of the SAM population

Fig. S10. GO enrichment analysis of genes associated with broomrape resistance

Fig. S11. Volcano plot of DEGs for broomrape resistance

Fig. S12. Ven diagram showing shared genes among analyses

Fig. S13. Comparison of genomic regions clusters associated with broomrape resistance across reference genomes.

Table S1. Summary of SAM population response to sunflower broomrape

Table S2. Summary table for the analysis of variance between the four main genetic groups among the SAM population in response to Yavor and Gadot broomrapes

Table S3. Summary table for the analysis of variance between Yavor and Gadot broomrapes

Table S4. Summary table for the analysis of variance of the field experiment

Table S5. Summary of phenotypic data collected in the field experiment for Yavor and Gadot broomrape populations

Table S6. SNPs and k-mers significantly associated with broomrape resistance and annotated genes in HA412OH, XRQ, PSC8 and LR1 genomes

Table S7. Significantly enriched GO terms of annotated genes associated with response to broomrape infection

Table S8. Annotated genes of significantly associated markers coincide Introgressed regions in HA412OH, XRQ, PSC8 and LR1 genomes

Table S8. Differentially expressed genes of infested and non-infested roots of a resistant sunflower cultivar 'EMEK3'

*Tables S5 – S8 Provided as an Table_data.xlsx document

### Supplementary figures:

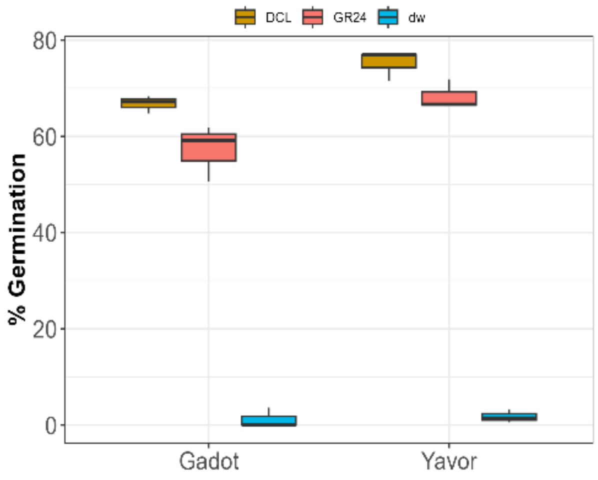

**Fig. S1.** Evaluation of germination rates among 'Yavor' and 'Gadot' broomrape seeds treated with $10^{-5}$M of the artificial germination stimulants GR24 (red), DCL (tan) and distilled water (blue) as control. Each box depicts a different treatment for each broomrape type, where the box represents the 1st and 3rd quartiles, horizontal line is the median, and whiskers represent the variance.

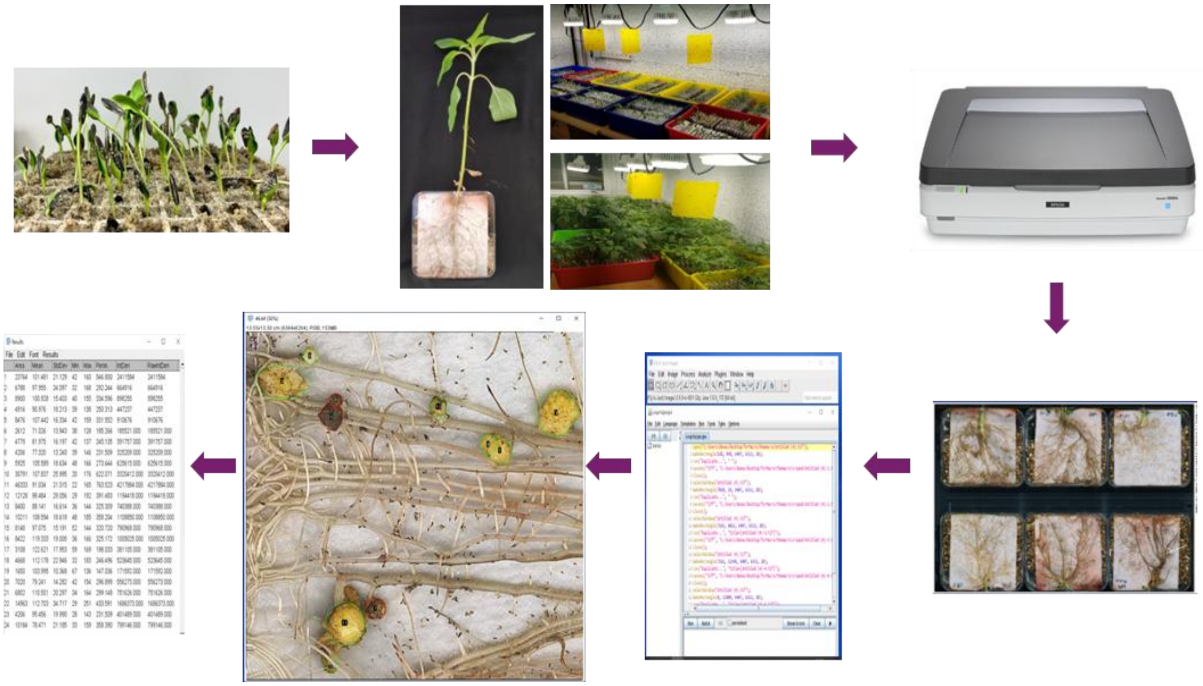

**Fig. S2.** High-throughput phenotyping for the response to broomrape infestation using the rhizotron platform and image analysis. Workflow from topleft: sunflower seeds are germinated in trays and transplanted to a squared petri dish. After 35 days, roots are scanned with high-resolution scanner and images are analyzed using dedicated macro in imageJ software.

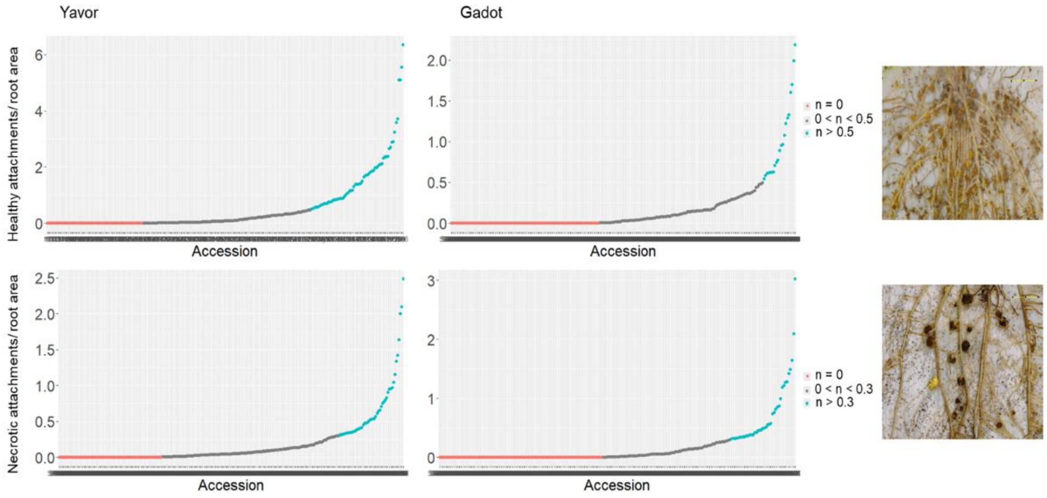

**Fig. S3.** Scatter plot of HealAD (top) and NecAD (bottom) measured in response infestation with ‘Yavor’ (left) and ‘Gadot’ (right) broomrapes. Accessions are sorted by their measured phenotype, where on the left are the accessions selected for the extreme low phenotypes (red) and on the right are the extreme high phenotypes (blue). The specific thresholds are indicated to the right of the plots. On the right are images illustrating the HealAD and NecAD phenotypes, respectively.

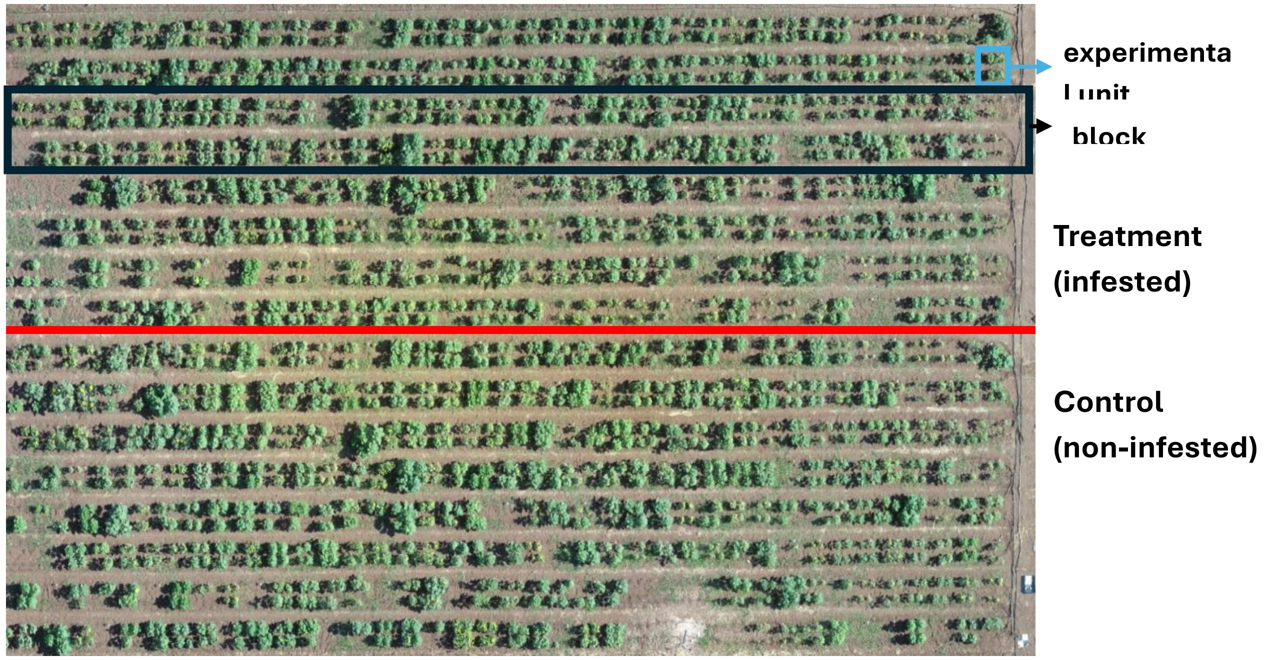

**Fig. S4.** A photo of the field experiment in "Gadash" farm taken from a drone. The experimental design includes two treatments (separation indicated with red line), four blocks for each treatment (black rectangle) and a plot for each accession in each block (blue rectangle).

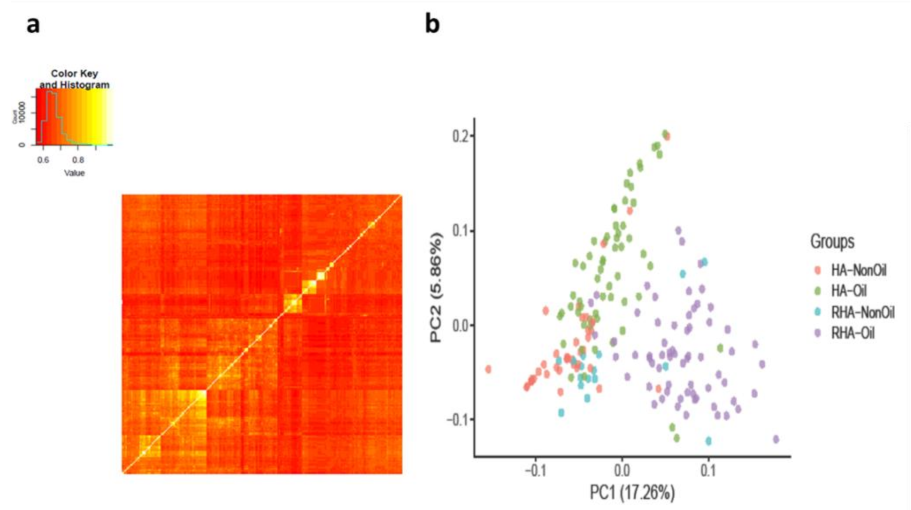

**Fig. S5.** Evaluating population structure and relatedness in the SAM population. (a) Kinship matrix for all accessions, where light color indicates higher relatedness, and (b) scatterplot for the two first PCs from a PCA over all SNP data. The four different genetic groups are indicated with different colors.

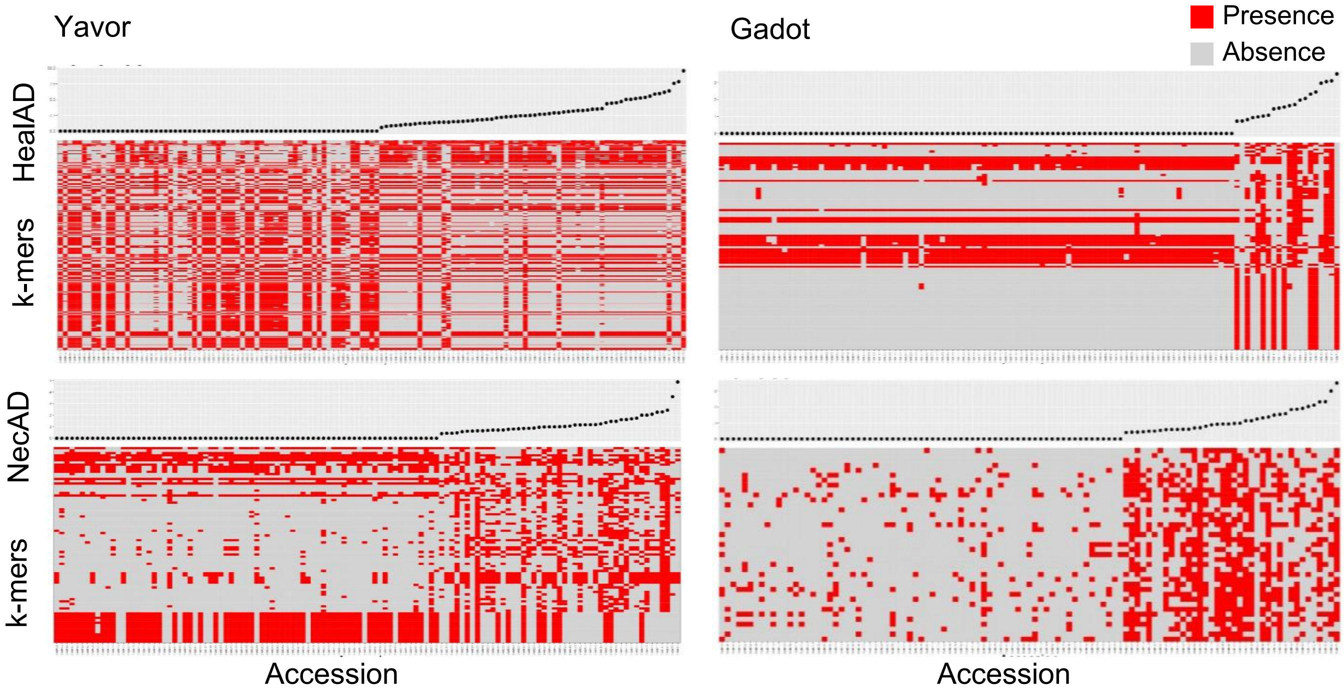

**Fig. S6.** Genomic composition of accessions with extreme phenotypes. At the top of each plot presented are the phenotypes score of each accession and at the bottom are the presence (red) or absence (grey) of significantly associated *k*-mers. The two upper plots correspond to HealAD, and the plots at the bottom correspond to NecAD phenotypes in response to infestation with 'Yavor' (left) and 'Gadot' (right) broomrapes.

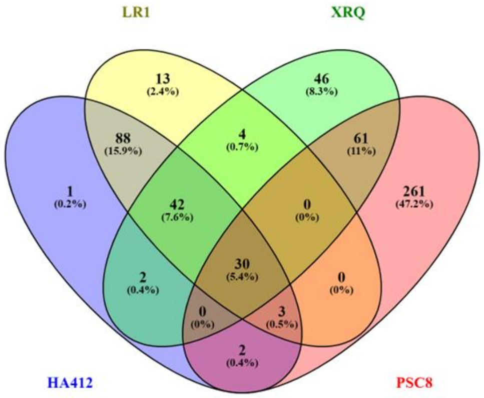

**Fig. S7.** A Venn diagram showing the number of shared *k*-mers among HA412OHv2, XRQv2, PSC8 and LR1 genomes.

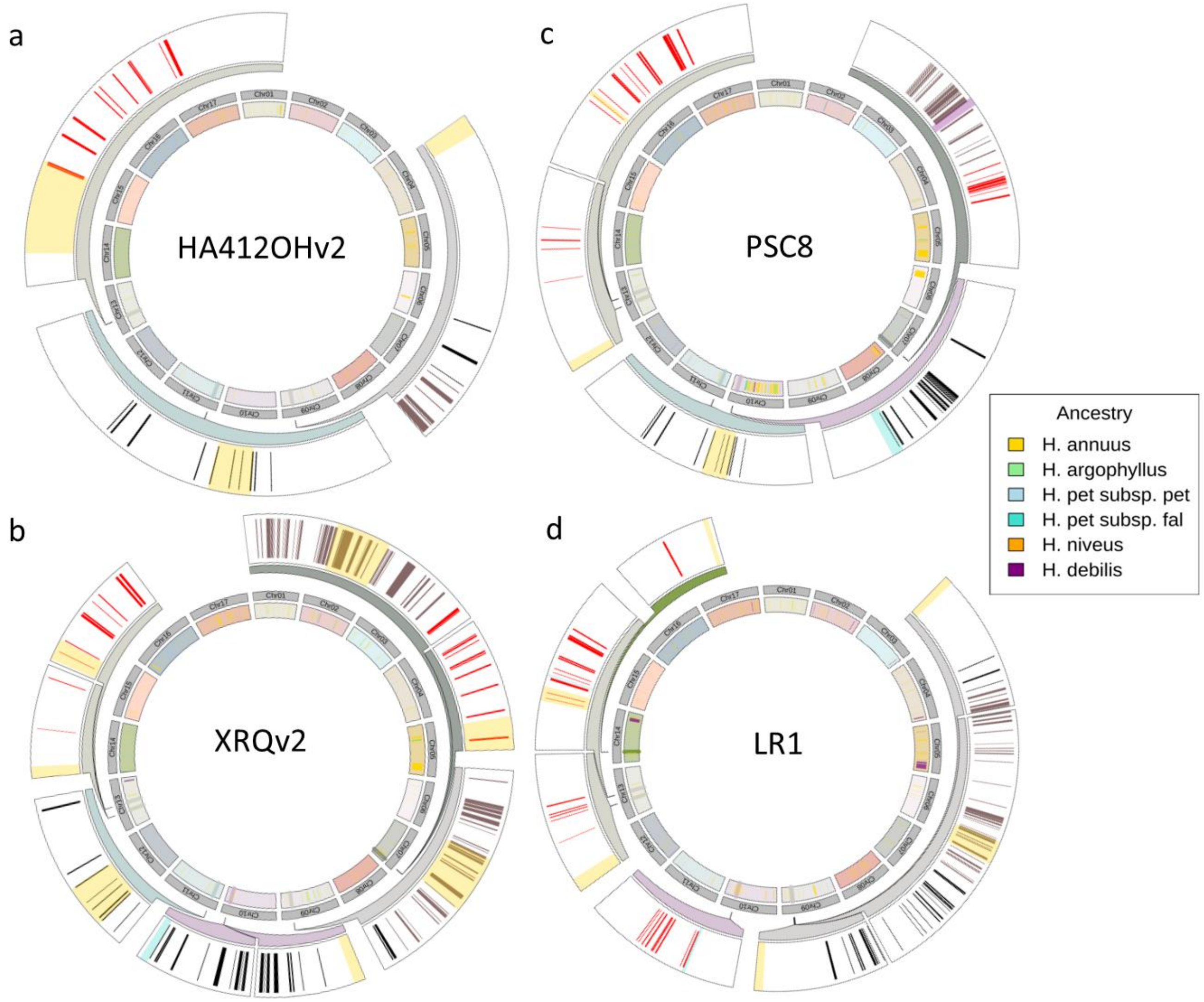

**Fig S8.** Circular plots showing the genes within ± 250kbp of associated genomic region based on SNPs (red bars) or *k*-mer (black bars) in the magnified outer circle. Overlaps with genomic regions that were identified as introgressions from wild sunflower species across all chromosomes in the inner circle (indicated with different colors). The four circular plots correspond to the HA412OHv2 (a) XRQv2 (b) PSC8 (c) and LR1 (d) genomes.

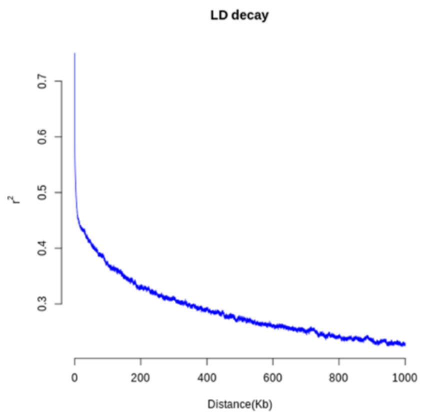

**Fig. S9.** Pattern of LD-decay in the SAM population.

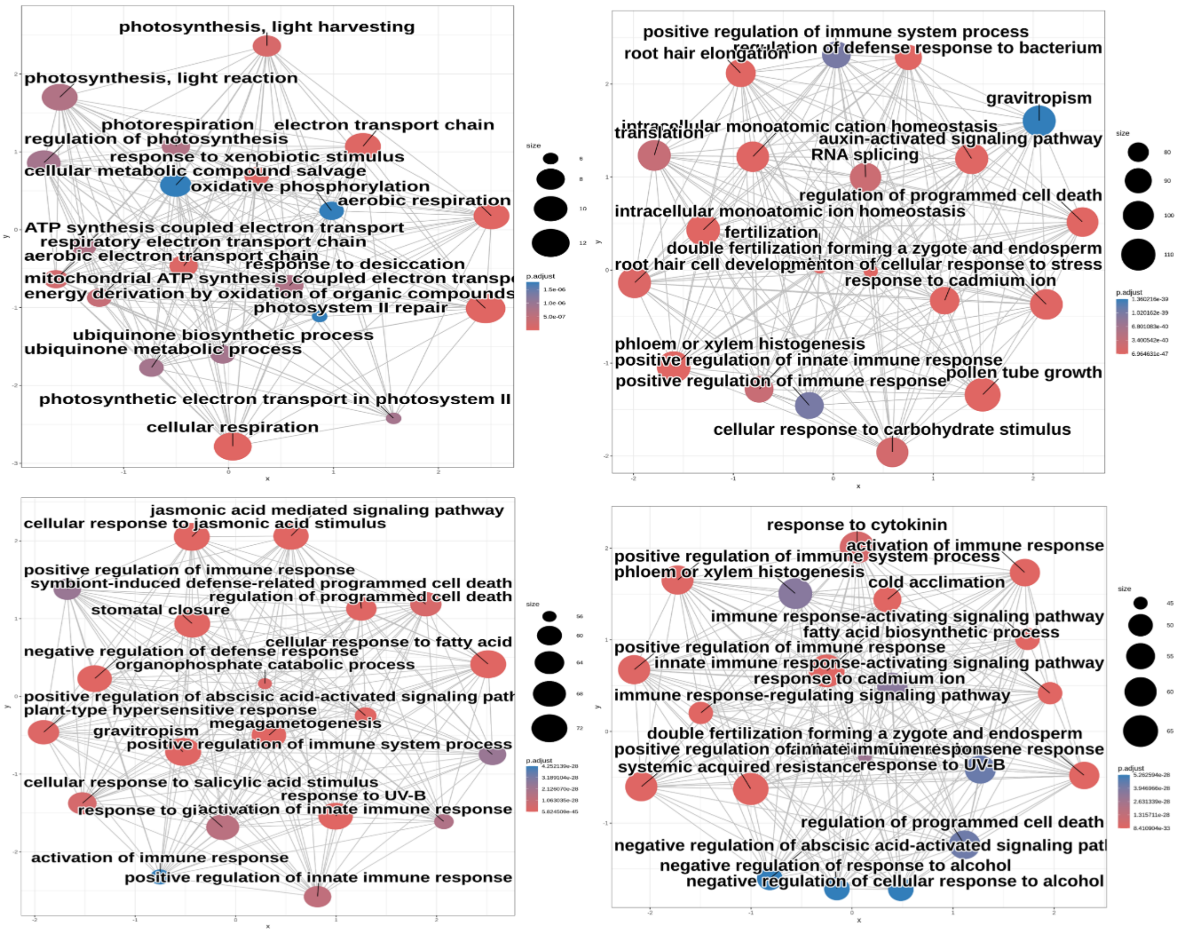

**Fig. S10.** Gene ontology (GO) enrichment pathway maps of annotated genes that are associated with HealAD (left) NecAD (right) in response to ‘Yavor’ race (top row), and annotated genes associated with HealAD (left) NecAD (right) in response to ‘Gadot’ broomrape (bottom row). In each plot, the size of the circle corresponds to the number of genes associated with the GO term and color (red-blue) correspond to p-values after FDR correction.

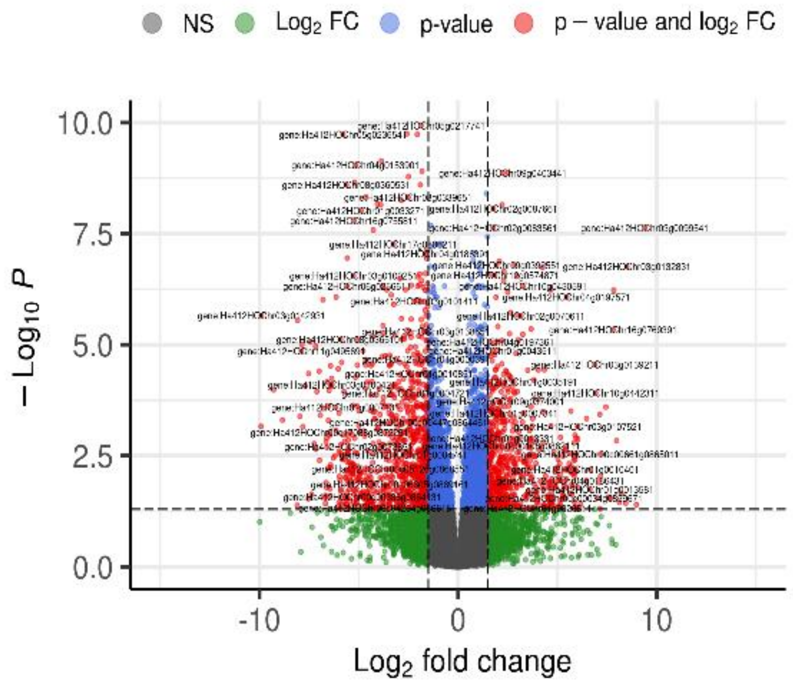

**Fig. S11.** Volcano plot of differentially expressed genes (DEGs) in the resistant hybrid variety 'EMEK3' five days post-infestation with and without 'Yavor' broomrape. Significantly differentially expressed genes are marked in red.

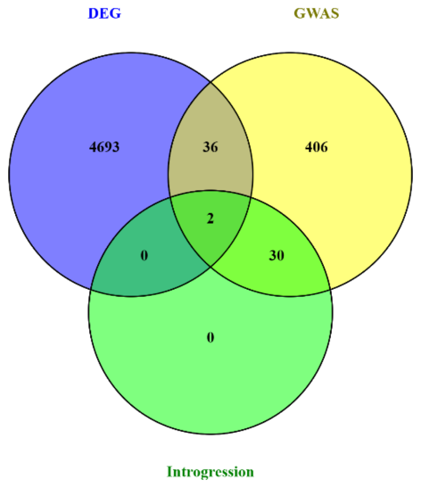

**Fig. S12.** A Venn diagram showing the number of shared genes between the GWAS analyses, DEGs in the RNA-Seq analyses, and introgressions identified in the HA412OHv2 genome.

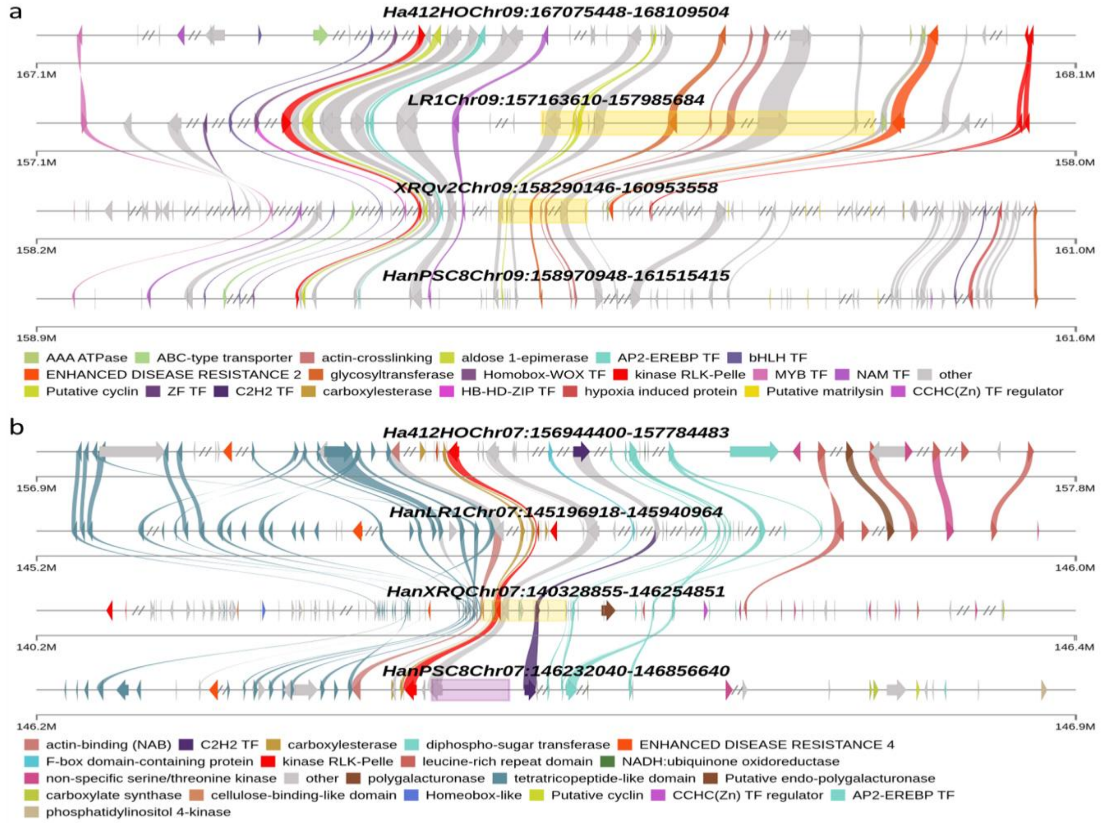

**Fig. S13.** Zoom in on genomic regions identified as clusters of genes mapped to the different reference genomes on chromosome 9 (top) and chromosome 7 (bottom). Candidate resistant genes are marked with colors and nearby introgressed regions from wild species are indicated with colored horizontal bar: wild *H. annuus* (yellow) or from *H. debilis* (purple).

### Supplementary tables:

**Table S1.** Summary of the response to ‘Yavor’ and ‘Gadot’ broomrapes.

| | Mean No. of attachments | % Healthy | % Necrotic | No. resistant accessions (attachments = 0) |
|---|---|---|---|---|
| **Yavor** | 43.05 | 73.7 | 26.3 | 66 |
| **Gadot** | 18.3 | 46.8 | 53.2 | 95 |

**Table S2.** Summary statistics of post-hoc analysis of the response to infestation with each broomrape race among the four breeding groups in the SAM population.

| Phenotype | Broomrape | Sunflower group | emmean | SE | asymp.LCL | asymp.UCL | post-hoc |
|---|---|---|---|---|---|---|---|
| HealAD | Yavor | HA_NonOil | 1.09925 | 0.15806 | 0.82928 | 1.45710 | A |
| | | HA_Oil | 0.61330 | 0.09568 | 0.45174 | 0.83265 | B |
| | | RHA_NonOil | 1.40745 | 0.27217 | 0.96345 | 2.05606 | A |
| | | RHA_Oil | 1.03341 | 0.12513 | 0.81509 | 1.31021 | A |
| | Gadot | HA_NonOil | 0.55652 | 0.12102 | 0.36340 | 0.85227 | A |
| | | HA_Oil | 0.24076 | 0.06086 | 0.14670 | 0.39514 | AB |
| | | RHA_NonOil | 0.55655 | 0.18649 | 0.28859 | 1.07330 | AB |
| | | RHA_Oil | 0.18493 | 0.05646 | 0.10165 | 0.33643 | B |
| NecAD | Yavor | HA_NonOil | 0.62312 | 0.11900 | 0.42856 | 0.90599 | A |
| | | HA_Oil | 0.25549 | 0.06175 | 0.15909 | 0.41030 | B |
| | | RHA_NonOil | 0.35201 | 0.13611 | 0.16498 | 0.75109 | AB |
| | | RHA_Oil | 0.21022 | 0.05644 | 0.12421 | 0.35579 | B |
| | Gadot | HA_NonOil | 0.52874 | 0.11796 | 0.34146 | 0.81873 | A |
| | | HA_Oil | 0.28291 | 0.06597 | 0.17912 | 0.44683 | AB |
| | | RHA_NonOil | 0.47615 | 0.17251 | 0.23408 | 0.96858 | AB |
| | | RHA_Oil | 0.15082 | 0.05099 | 0.07774 | 0.29259 | B |

**Table S3.** Summary table for the analysis of variance between 'Yavor' and 'Gadot' populations for total, healthy and necrotic tubercle normalized to root area.

| | | Df | Sum Sq | Mean Sq | F value | Pr(>F) | |
|---|---|---|---|---|---|---|---|
| TotAD | broomrape population | 1 | 12.5 | 12.486 | 12.88 | 3.81E-04 | *** |
| | Residuals | 336 | 325.6 | 0.969 | | | |
| HealAD | broomrape population | 1 | 12.5 | 12.503 | 22 | 3.97E-06 | *** |
| | Residuals | 336 | 190.9 | 0.568 | | | |
| NecAD | broomrape population | 1 | 0 | 0.00001 | 0 | 0.995 | |
| | Residuals | 336 | 49.53 | 0.14741 | | | |

**Table S4.** Summary table for the analysis of variance for the different traits measured in each field trial.

| Broomrape | Source of variance | Df | F value | | | | | |
|---|---|---|---|---|---|---|---|---|
| | | | Plant height | Stem diameter | Main flower diameter | Days to budding | Days to R55 | Days to anthesis |
| **Yavor** | Treatment | 1 | 0.30 | 38.9307*** | 1.295 | 45.5915*** | 1.8544 | 2.6315. |
| | Accession | 76 | 20.1361*** | 13.0912*** | 21.6034*** | 11.8877*** | 34.7335*** | 29.2072*** |
| | Treatment*Accession | 76 | 1.18 | 0.86 | 0.9778 | 1.2962. | 1.05 | 1.1907 |
| **Gadot** | Treatment | 1 | 0.09 | 5.4086* | 1.2253 | 1.2527 | 0.0902 | 3.2372. |
| | Accession | 76 | 10.5038*** | 3.5477*** | 4.0276*** | 1.5465** | 13.0982*** | 14.3707*** |
| | Treatment*Accession | 76 | 1.1538 | 0.9864 | 1.1584 | 0.8542 | 1.5953** | 1.477** |
| | Signif. codes: 0 '***' 0.001 '**' 0.01 '*' 0.05 '.' | | | | | | | |
